# IKKα and IKKβ serve as kinase-independent scaffolds that prevent proteasomal degradation of NEMO

**DOI:** 10.64898/2026.09.10.750631

**Authors:** Kathryn Balmanno, Jack A. Prescott, Nadav Myers, Eleanor J. Griffiths, Yosef Shaul, Simon J. Cook

## Abstract

NF-κB essential modulator (NEMO) is the critical scaffold protein of the canonical I kappa B kinase (IKK) complex. Whilst studying the contribution of IKKα and IKKβ to NF-κB signalling we noted that NEMO protein levels were reduced in IKK knockout (KO) HCT116 cells. Loss of NEMO was more pronounced in IKKα KO cells than IKKβ KO, whilst IKKα+IKKβ double knockout (DKO) cells exhibited a 90% reduction in NEMO abundance. In contrast, the abundance of IKKs was unaffected by NEMO KO, although this abolished canonical NF-κB activation. Loss of IKKs had no effect on NEMO mRNA levels but IKK DKO cells exhibited accelerated NEMO protein turnover that was reversed by the proteasome inhibitor, MG132, but not the NEDDylation inhibitor, MLN4924. TAK-243, a selective inhibitor of UBA1, the E1 ubiquitin-activating enzyme, increased basal NEMO levels in DKO cells but did not prevent NEMO turnover suggesting both ubiquitylation-dependent and independent pathways for degradation. Bioinformatic analysis identified substantial regions of intrinsic disorder within the N- and C-termini of NEMO. NEMO was rapidly degraded by isolated 20S proteasomes *in vitro* and this was inhibited by interaction with IKKα or IKKβ. Finally, re-expression of IKKα increased NEMO expression in IKK DKO cells and this required the C-terminal NEMO binding domain (NBD) but not a functional kinase domain. Thus the IKKs act as critical kinase-independent scaffolds that stabilise NEMO; loss of these interactions results in NEMO degradation via the proteasome. These results may be relevant to strategies for inhibiting NF-kB signalling by disrupting NEMO:IKK interactions.

## INTRODUCTION

The canonical Inhibitor of kappa B kinase (IKK) complex, the key regulator of the NF-κB signalling [1], is minimally composed of three components: the homologous IKKs (IKKα and IKKβ), and a regulatory scaffold protein, NF-κB essential modulator (NEMO, also known as IKKγ) [2–4]. Although NEMO is catalytically inactive, NEMO-null cell lines fail to activate the canonical NF-κB pathway in response to almost all pro-inflammatory, immune regulatory and pro-survival stimuli [5] and NEMO-deficient mice die early in embryogenesis due to substantial liver apoptosis [6]. Point mutations and deletions in the *IKBKG* gene, encoding NEMO, cause a number of X-linked genetic disorders, including incontinentia pigmenti, a severe form of dermatitis [5, 7], and anhydrotic ectodermal dysplasia with immunodeficiency (EDA-ID) [8], characterised by severe bacterial infections of the respiratory, gastrointestinal tracts, bones, soft tissues and skin. These studies highlight the fundamental importance of NEMO as a scaffold in facilitating activation of the canonical IKK complex and NF-κB signalling.

NEMO is a 48 kilodalton multi-domain polypeptide [2] that forms an elongated, parallel intermolecular coiled coil [2,9,10]. NEMO contains two coiled coil (CC) domains, CC1 and CC2, two small α-helical regions (αH1, αH2) located either side of the CC1 domain, a leucine zipper (LZ), a proline rich region (PRR) and a zinc-finger domain (ZNF) at the extreme C-terminus. In addition, parts of CC2 and LZ, together with a conserved ubiquitin binding sequence (amino acids 285-315), make up the so-called ‘ubiquitin binding in ABIN and NEMO’ (UBAN) domain [11]. The ZNF domain of NEMO is thought to mediate interactions with polyubiquitin chains [12,13], in addition to IκBα [14]. A region spanning the αH1 and CC1 domains of NEMO, minimally defined as aa 44-111, mediates interaction with IKKα and IKKβ and is known as the IKK-binding domain (IBD) [15,16]. In turn, IKK binding to NEMO is mediated by a short C-terminal sequence (LDWSWL) in IKKα and IKKβ known as the NEMO-binding domain (NBD), which is required for IKK activation [15].

The precise *in vivo* composition of the IKK complex and the stimulus-dependency of this composition remain a matter of debate. Biochemical assays with purified IKK complexes suggest the minimal canonical IKK complex is composed of an IKKα:IKKβ heterodimer bound to a dimer of NEMO subunits [17,18]. Full-length NEMO has been described as existing in a dimeric, trimeric or tetrameric state [19], whereas hydrodynamic studies suggest that apo-NEMO (in the absence of IKK) is a dimer in weak equilibrium with a tetrameric assembly, and that interaction with IKK is mutually exclusive with tetramerization [20]. A region spanning the αH1 and CC1 domains that overlaps the IKK binding domain has been implicated in NEMO dimerization [9], while a region spanning CC2 and the LZ has been described as a minimal oligomerization domain [21]. Mutations in this domain greatly reduce NF-κB activation, suggesting that NEMO oligomerisation is important for activation of the NF-κB pathway [16]. A higher order NEMO lattice extending from 400 nm to over 1 μm has been observed in cells, and was speculated to be composed of NEMO dimers linked through dimeric IKK subunits and polyubiquitin chains [22].

We recently used CRISPR/Cas9 gene editing to generate IKKα KO or IKKβ KO HCT116 and SW620 colorectal cancer cell lines; this unexpectedly revealed that IKKα was more important than IKKβ for canonical activation of NF-κB by TNFα and IL-1 [23]. Here we show that loss of IKKα and IKKβ (IKK DKO) results in a striking 90% reduction in the steady state level of NEMO protein. Our analysis suggests NEMO is degraded by the proteasome in the absence of IKKs, which act as obligate scaffolds to maintain NEMO stability, independent of their kinase activity. These results may be relevant to the role of NEMO as a signalling hub and therapeutic strategies to inhibit NF-κB.

## RESULTS

### NEMO protein levels are reduced following knock out or knock down of IKKs

We previously generated WT, IKKα KO, IKKβ KO and IKKα/β DKO HCT116 cell lines by CRISPR/Cas9 gene editing [23]. Whilst studying the response of these cells to TNFα, we noted a reduction in NEMO protein abundance in IKK KO cells. Prolonged (24-48hrs) stimulation with TNFα had little effect on NEMO protein levels in WT cells (Figure 1A). Loss of IKKα caused a strong reduction in basal NEMO levels and revealed a modest, TNFα-dependent reduction in NEMO. Loss of IKKβ had little effect on basal NEMO levels but TNFα stimulation still resulted in reduction in NEMO. Finally, basal NEMO levels were substantially reduced in IKKα/β DKO cells regardless of TNFα-stimulation (Figure 1A).

**Figure 1.**
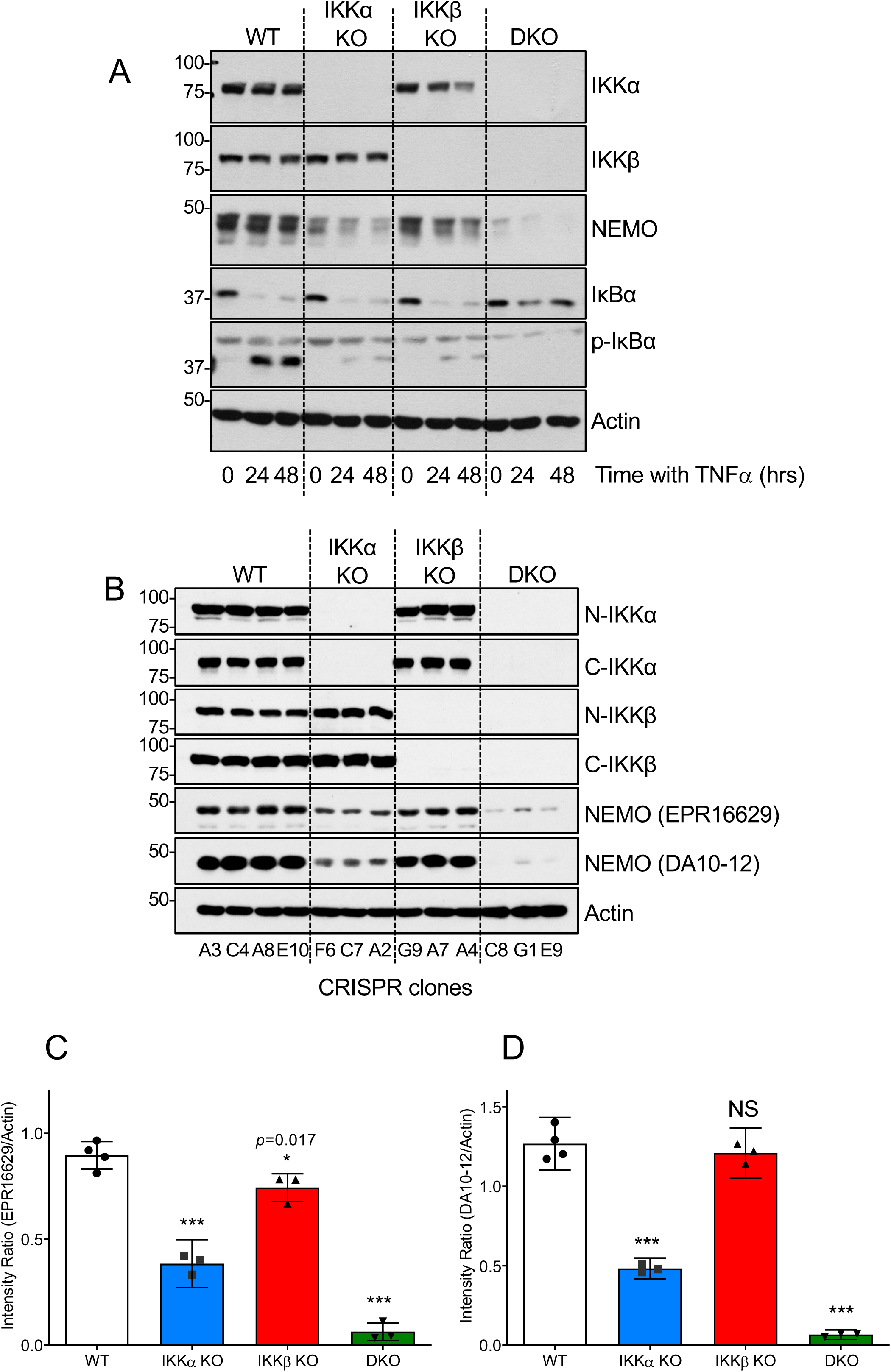
IKK knockout HCT116 cells exhibit a substantial reduction in NEMO protein abundance. **(A)** Single-cell derived WT (clone A3), IKKα KO (clone F6), IKKβ KO (clone G9) and IKKα/β DKO (clone C8) HCT116 cells were seeded in normal growth medium for 24 hours prior to treatment with 10 ngml^-1^ TNFα for 24 or 48 hours. Whole lysates were fractionated by SDS-PAGE and Western blotted with the indicated antibodies. **(B)** Whole-cell lysates generated from the indicated WT, IKKα KO, IKKβ KO and IKKα/β DKO HCT116 clones were fractionated by SDS-PAGE and Western blotted with the indicated antibodies. Data are from a single experiment representative of at least two others giving similar results. **(C and D)** Densitometry plots for western blot data shown from the experimental series described in (B). Intensities of bands were calculated using ImageJ, verifying for non-saturation and subtracting background. Intensity ratios after normalising against Actin loading control were plotted. Data in (C) corresponds to Western blot data using NEMO antibody EPR126229 (Abcam). Data in (D) corresponds to Western blot data using NEMO antibody DA10-12 (Cell Signalling Technology). Data points correspond to individual clones in (B) and represent mean ± SD. Statistical testing performed using one-way ANOVA between WT and KO samples, with Dunnett’s multiple comparisons test (*** =p<0.001).

Focusing on basal, unstimulated NEMO levels we compared four different WT clones with three different KO clones for each genotype by western blot using two different monoclonal antibodies against NEMO, so any apparent changes in abundance were unlikely to be due to IKK loss impacting a specific NEMO epitope. These results confirmed a reduction in basal NEMO levels (Figure 1B). Quantification of blots from multiple experiments with both NEMO antibodies demonstrated that IKKα knockout resulted in a significant 50-60% decrease in basal NEMO abundance, while IKKα/IKKβ DKO cells exhibited a striking >90% decrease in NEMO protein (Figure 1C & 1D); loss of IKKβ alone caused only a modest reduction in NEMO protein. We also employed reverse-transcription quantitative PCR (RT-qPCR) to determine if the reduction in NEMO expression in IKK KO cells was the result of decreased mRNA levels (Supplementary Figure 1A). Single or double IKK KO showed no significant reduction in NEMO mRNA levels. In contrast to the levels of NEMO protein, IKKα/β DKO cells actually exhibited a small increase in NEMO mRNA expression relative to WT, which might reflect an attempt by the cells to adapt and restore NEMO protein levels; however, this difference did not reach significance for the three DKO clones examined.

Interpreting protein abundance in knockout cells can be complicated by long-term adaptation, so we also employed transient siRNA-mediated knockdown of IKK subunits in HCT116 cells (Supplementary Figure 1B). IKKα knockdown caused a greater reduction in NEMO expression than IKKβ knockdown, while knockdown of both caused a striking reduction in NEMO abundance, consistent with results in the IKK KO cells. Loss of NEMO was also observed in other cell lines; for example, IKKα KO SW620 cells [23] exhibited a 50% reduction in basal NEMO levels and this was further reduced, to 75-80%, by siRNA-mediated IKKβ knockdown (Supplementary Figure 2A & 2B).

Collectively, these data demonstrated that IKKs are necessary to maintain expression of NEMO protein; that IKKα appears to be of greater importance than IKKβ; and the loss of NEMO protein is not due to a reduction in NEMO mRNA.

### Loss of NEMO expression has no impact on IKKα or IKKβ abundance

To address the reciprocal question - whether loss of NEMO influences IKK abundance - we used CRISPR/Cas9 gene editing to generate multiple NEMO KO clones. Western blot analysis indicated that loss of NEMO with either of two different guides had no effect on the basal level of IKKα or IKKβ (Figure 2A). When these cells were stimulated with TNFα the parental (WT) and CRISPR control cells (C) responded as expected with rapid phosphorylation of IKKs, degradation IκBα, and phosphorylation of p65; these responses were lost in NEMO KO clones but the level of IKKα or IKKβ was unaffected by NEMO loss (Figure 2B). Finally, TNFα-dependent activation of a multimerised NF-κB:Luc reporter was completely lost in the NEMO KO cells (Figure 2C), consistent with the role of NEMO as a critical scaffold for activation of the IKKs. Thus, whilst IKKs were necessary to maintain expression of NEMO protein (Figure 1), NEMO was not required to maintain expression of IKKα or IKKβ, though it was of course required for canonical TNFα-stimulated activation of the IKKs.

**Figure 2.**
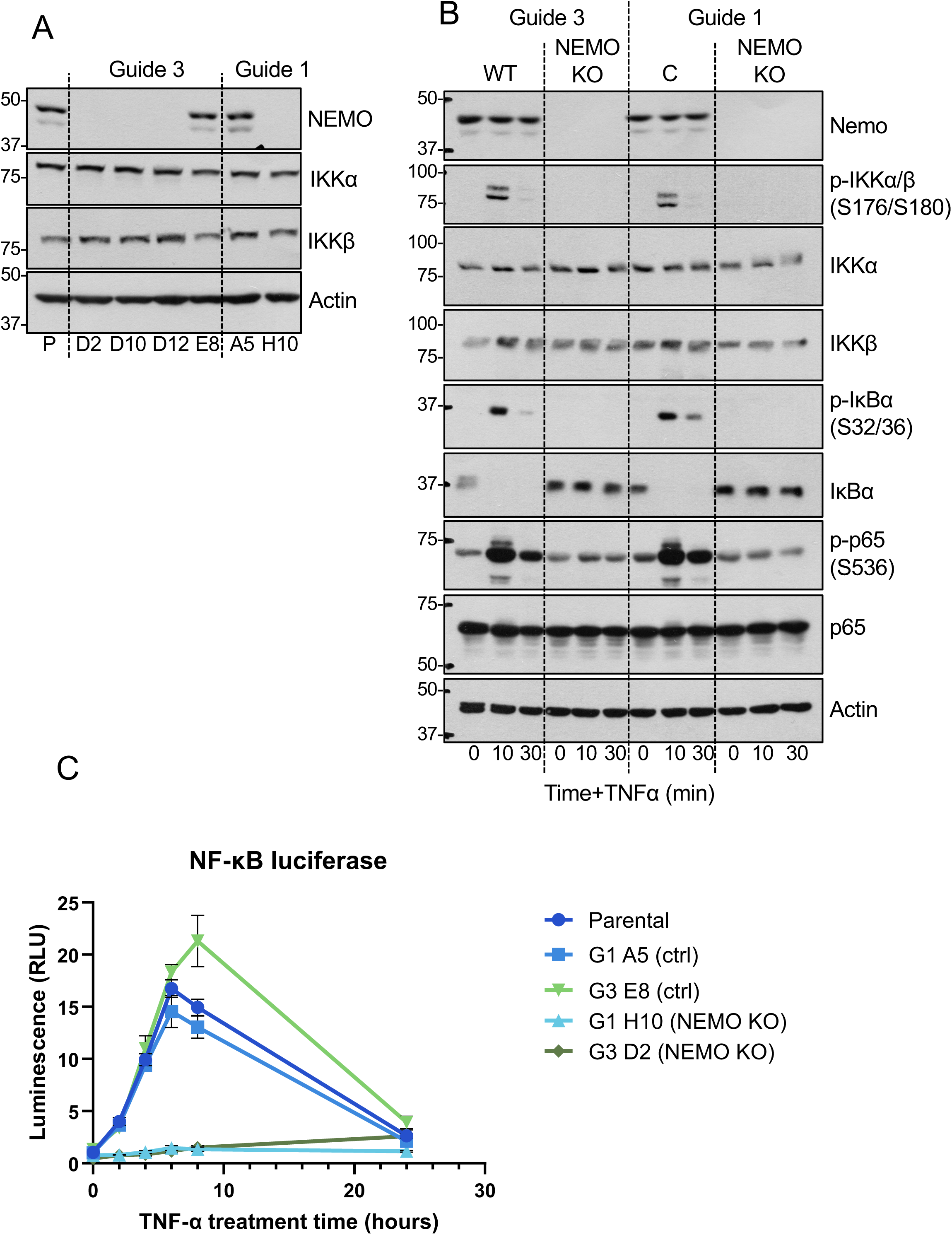
NEMO knockout does not reduce expression of IKKα or IKKβ but abolishes their activation. **(A)** Whole-cell lysates were generated from the indicated single-cell derived CRISPR/Cas9 NEMO KO HCT116 cell clones (or controls, E8, A5), fractionated by SDS-PAGE and Western blotted with the indicated antibodies. **(B)** Parental (WT), CRISPR Control (C) or NEMO KO HCT116 cells (clones D2 and H10) were stimulated with 10 ngml^-1^ TNFα for 10 or 30 mins. Whole-cell lysates were fractionated and Western blotted with the indicated antibodies. A single experiment is shown; identical results were obtained in two additional experiments. **(C)** Parental, CRISPR Control (Ctrl) or NEMO KO HCT116 cells were transiently transfected with pGL4.32 luc2P/NF-κB-RE firefly luciferase and pCMV renilla luciferase. 24 hours later, cells were stimulated with 10 ngml^-1^ TNFα for 2-24 hours. Firefly luciferase luminescence was normalised relative to renilla luciferase luminescence. Results are mean ± SD of three experiments performed in cell culture triplicate.

### NEMO undergoes proteasome-dependent turnover in the absence of IKKs

To investigate mechanism underlying the loss of NEMO protein we used chemical inhibitors of the main protein degradation systems in cells – the ubiquitin proteasome system (UPS) and the Lysosome. Representative experiments are shown in Supplementary Figure 3A and 3B and are quantified in Supplementary Figure 3>C. Treatment of IKKα/β DKO cells with MG132 (a proteasome inhibitor) for 16 hours caused only a modest increase in the low basal level of NEMO that was not statistically significant (Supplementary Figure 3A-C). The Cullin-RING Ligases (CRLs) are the largest family of E3s, with more than 200 distinct E3 ligase complexes made up from different cullin scaffolds [24,25]. CRLs all require NEDDylation of the cullin component for activity and this is catalysed by a cascade of NEDDylating enzymes, including a dedicated E1 NEDD8-activating enzyme [26, 27]. Treatment of IKKα/β DKO cells with MLN4924, a potent and highly selective inhibitor of the E1 NEDD8-activating enzyme [28], also caused a modest increase in the low basal level of NEMO but this was also not statistically significant (Supplementary Figure 3A-C). Chloroquine (which disrupts lysosome function) failed to increase basal NEMO levels in IKKα/β DKO cells, but did increase basal LC3B levels indicating that the drug was effective (Supplementary Figure 3A-C). The modest increase in NEMO levels with MG132 or MLN4924 in the IKK DKO cells might indicate that the rates of transcription and/or translation of NEMO are low. It is conceivable that a longer treatment might provide a greater recovery but this was not possible with due to toxicity associated with long term treatment with these inhibitors.

**Figure 3.**
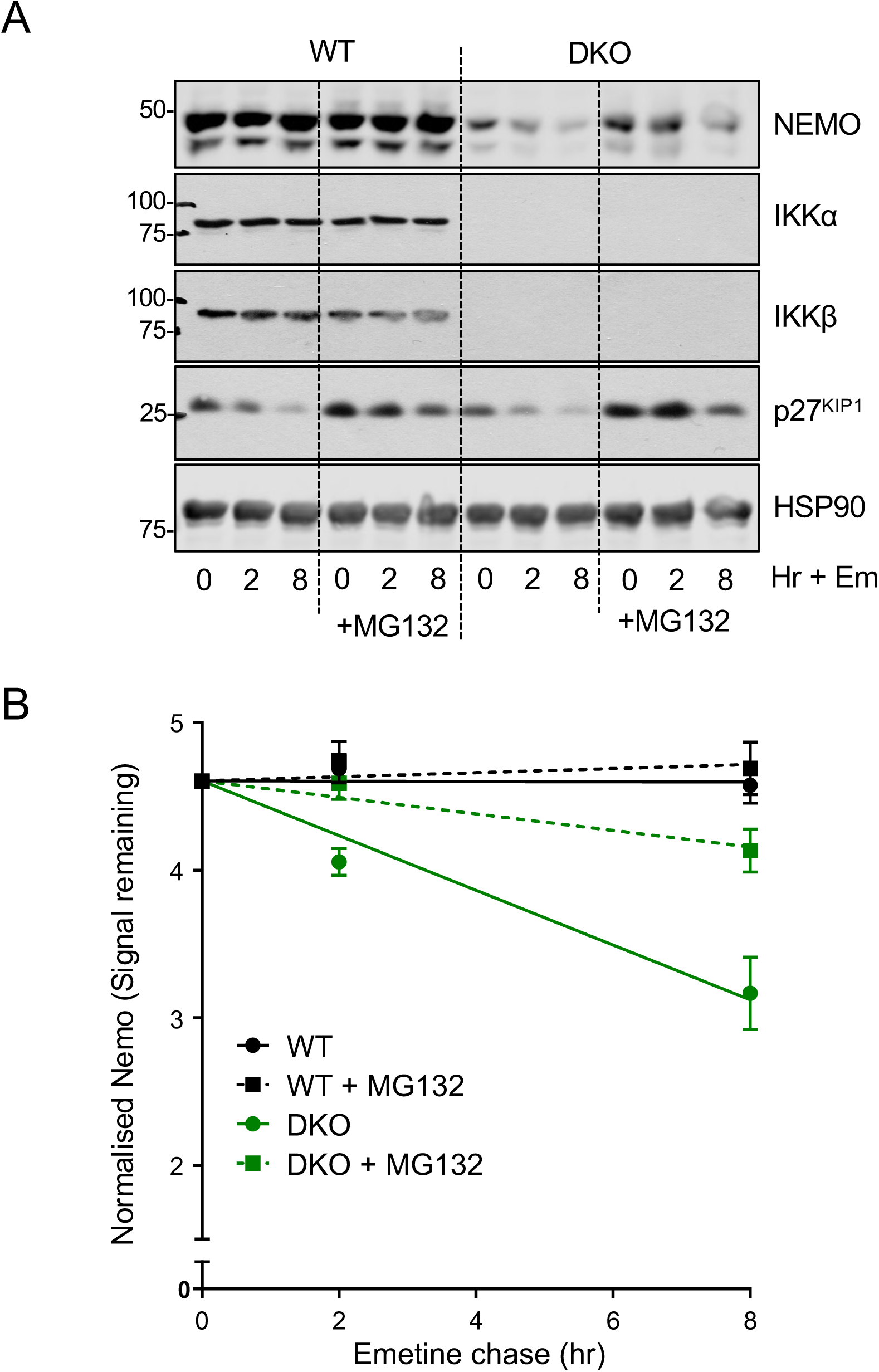
NEMO undergoes accelerated proteasome-dependent turnover in IKK DKO cells compared to WT cells. **(A)** WT or IKKα/β DKO HCT116 cells were pre-treated for 30 minutes with 10μM MG132 or vehicle before addition of 10μM emetine for 0, 2 and 8 hours. Whole cell lysates were fractionated by SDS-PAGE and Western blots carried out using the indicated antibodies. Data are from a single representative experiment. **(B)** Data from n=3 experiments in (A) were quantified by LiCor imaging and NEMO band intensity relative to HSP90 loading control calculated. Results are NEMO signal remaining compared to the ‘no emetine’ control and are mean ± SD of three independent experiments.

As an alternative we assessed the half-life of NEMO in WT and IKKα/β DKO cells by performing shorter duration ‘chase’ experiments with emetine dihydrochlorohydrate (hereafter Emetine), an alkaloid that blocks protein synthesis in eukaryotic cells by binding to the 40S ribosomal subunit and inhibiting the translocation step of the elongation cycle [29]. p27^KIP1^, a protein that is degraded by the proteasome, was used as a positive control; 2 or 8 hours of Emetine caused time-dependent loss of p27^KIP1^ that was largely prevented by inclusion of MG132 (Figure 3A). As seen previously, IKKα/β DKO cells under steady state conditions exhibited considerably less NEMO protein than WT cells (Figure 3A). Upon treatment with Emetine, NEMO was degraded at a much faster rate in DKO cells than in WT cells and this accelerated decay was strongly inhibited by MG132 (Figure 3A & 3B). Thus, the accelerated turnover of NEMO protein in IKKα/β DKO cells was dependent upon the proteasome. The rapid degradation of p27^KIP1^ in the presence of Emetine was also prevented by inclusion of MLN4924 (Figure 4A), consistent with p27^KIP1^ being degraded by Cullin-1-dependent SCF^Skp2^ E3 ligase complex [30]. In contrast, MLN4924 did not prevent the degradation of NEMO (Figure 4A and 4B).

**Figure 4.**
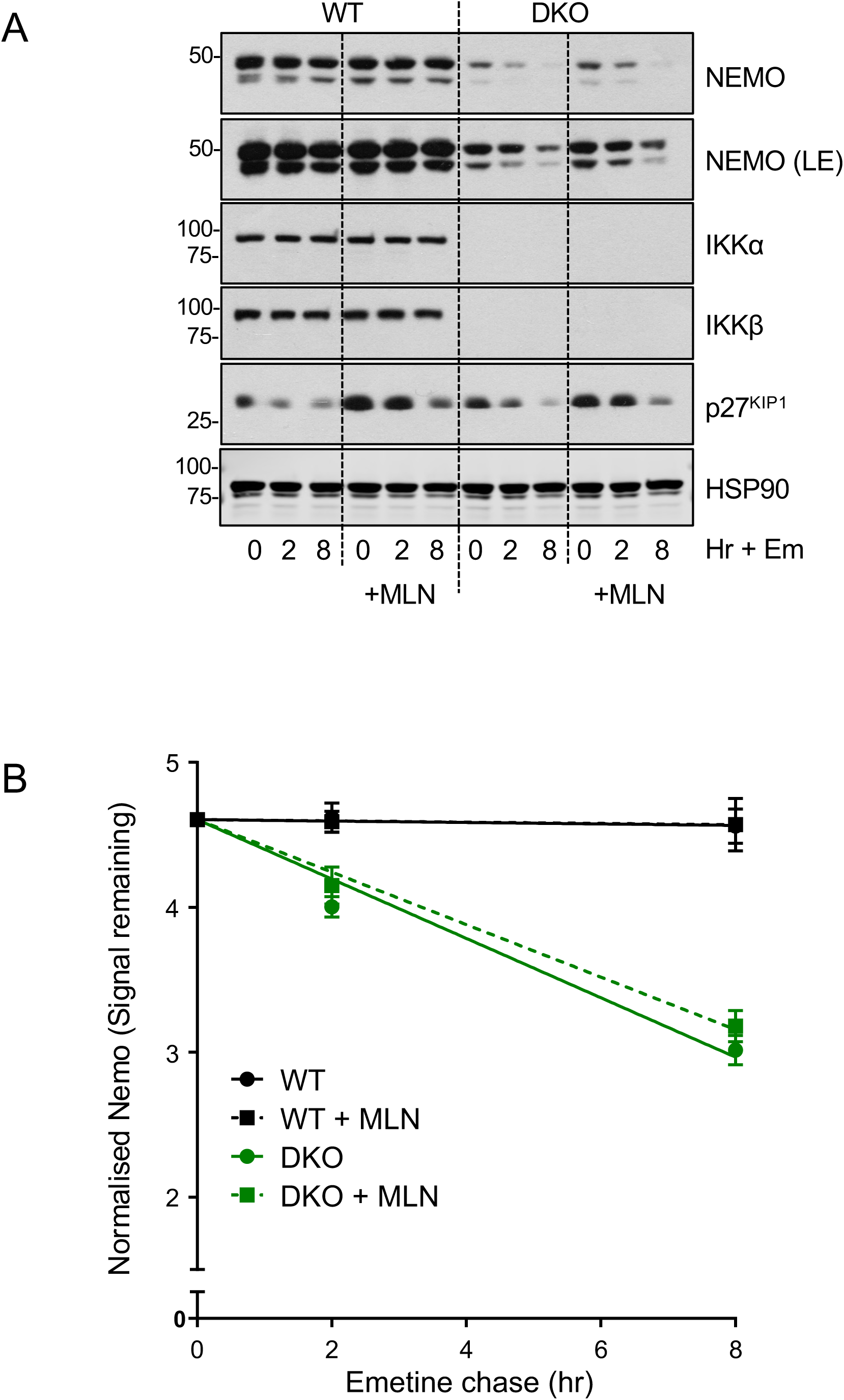
NEMO turnover in IKK DKO cells is Cullin-independent. **(A)** WT or IKKα/β DKO HCT116 cells were pre-treated for 30 minutes with 300nM MLN4924 before addition of 10μM emetine for 0, 2 and 8 hours. Whole cell lysates were fractionated by SDS-PAGE and Western blots carried out using the indicated antibodies. Data are from a single representative experiment. **(B)** Data from n=3 experiments in (A) were quantified by LiCor imaging and NEMO band intensity relative to HSP90 loading control calculated. Results are NEMO signal remaining compared to the ‘no emetine’ control and are mean ± SD of three independent experiments.

Together these results demonstrated that NEMO degradation is greatly accelerated in IKKα/β DKO cells compared to WT cells, this degradation is dependent on the proteasome but is independent of CRL E3 ligases.

### The UBA1 ubiquitin-activating enzyme inhibitor TAK-243 suggests two pathways for NEMO degradation in IKK DKO cells

Since CRL E3 ligases were not implicated in NEMO degradation in IKK DKO cells we used TAK-243, a potent and highly selective inhibitor of UBA1, the E1 ubiquitin-activating enzyme that is responsible for the bulk of cellular ubiquitylation [31]. Emetine chase experiments revealed that TAK-243 had no effect on the accelerated half-life of NEMO in IKK DKO cells, whereas the rapid degradation of p27^KIP1^ in the presence of Emetine was clearly slowed by TAK-243 (Figure 5A & 5B). However, we noted that an 8 hour TAK-243 treatment did increase the basal level of NEMO prior to the addition of Emetine (Figure 5A). Prompted by this observation we performed a time course study and observed that TAK-243 caused a gradual accumulation of both NEMO and p27^KIP1^ in IKK DKO cells (Figure 6A & 6B). These results suggested the presence of two pathways for NEMO degradation in IKK DKO cells; inhibition of the core ubiquitylation pathway by TAK-243 increased basal NEMO levels, indicating a ubiquitylation-dependent pathway, but NEMO could still be degraded in the presence of Emetine in TAK-243-treated cells.

**Figure 5.**
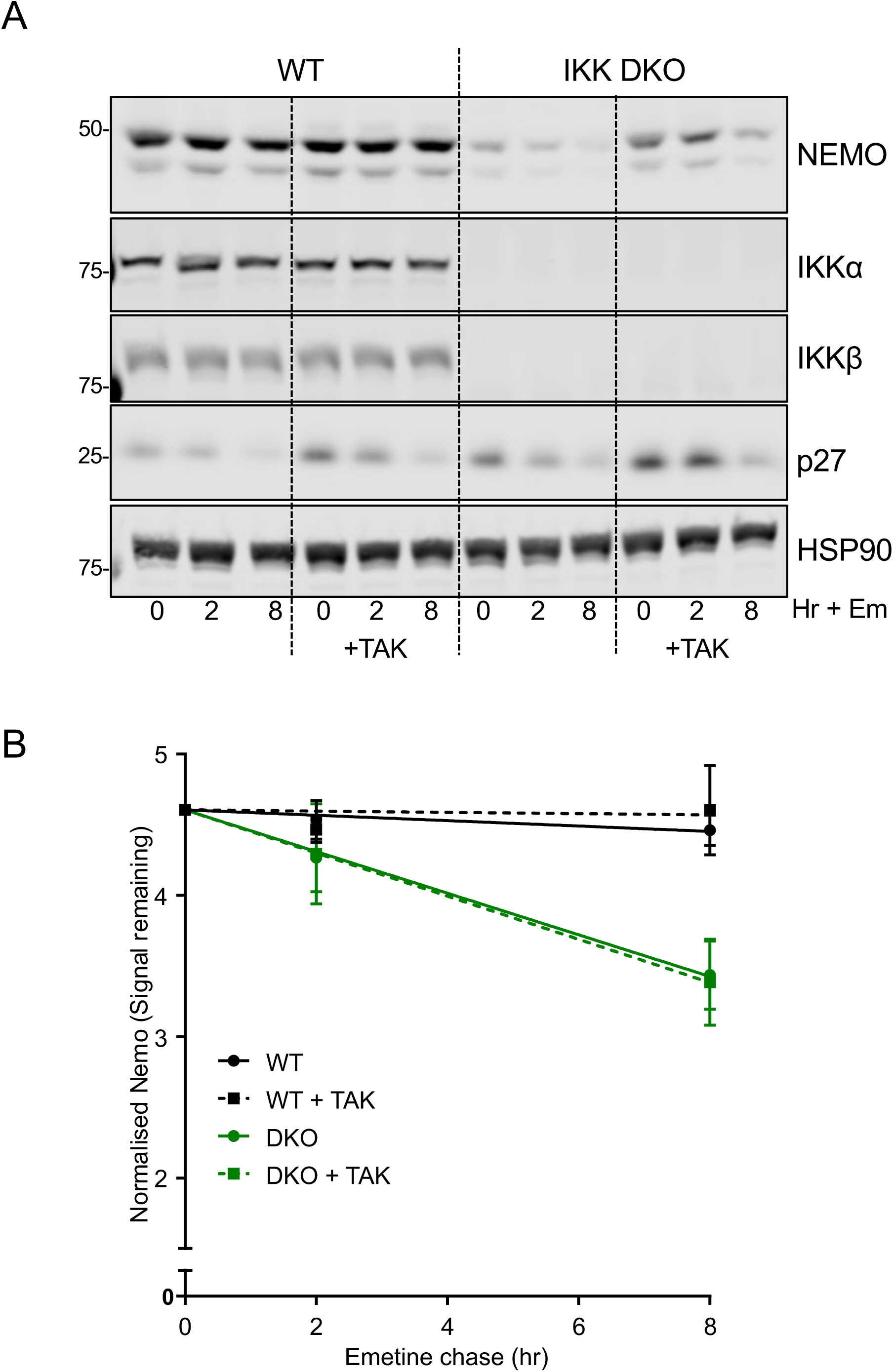
NEMO turnover in IKK DKO cells is not inhibited by TAK-243, a UBA1 E1 ubiquitin-activating enzyme inhibitor. **(A)** Where required IKK WT or IKKα/β DKO HCT116 cells were pre-treated for 30 minutes with 100nM TAK243 before addition of 10μM emetine for 0, 2 and 8 hours such that TAK243 treated cells were exposed to inhibitor for the same overall time. Whole cell lysates were fractionated by SDS-PAGE and Western blots carried out using the indicated antibodies. Data are from a single representative experiment. **(B)** Data from n=3 experiments in (A) were quantified by LiCor imaging and NEMO band intensity relative to HSP90 loading control calculated. Results are NEMO signal remaining compared to the ‘no emetine’ control and are mean ± SD of three independent experiments.

**Figure 6.**
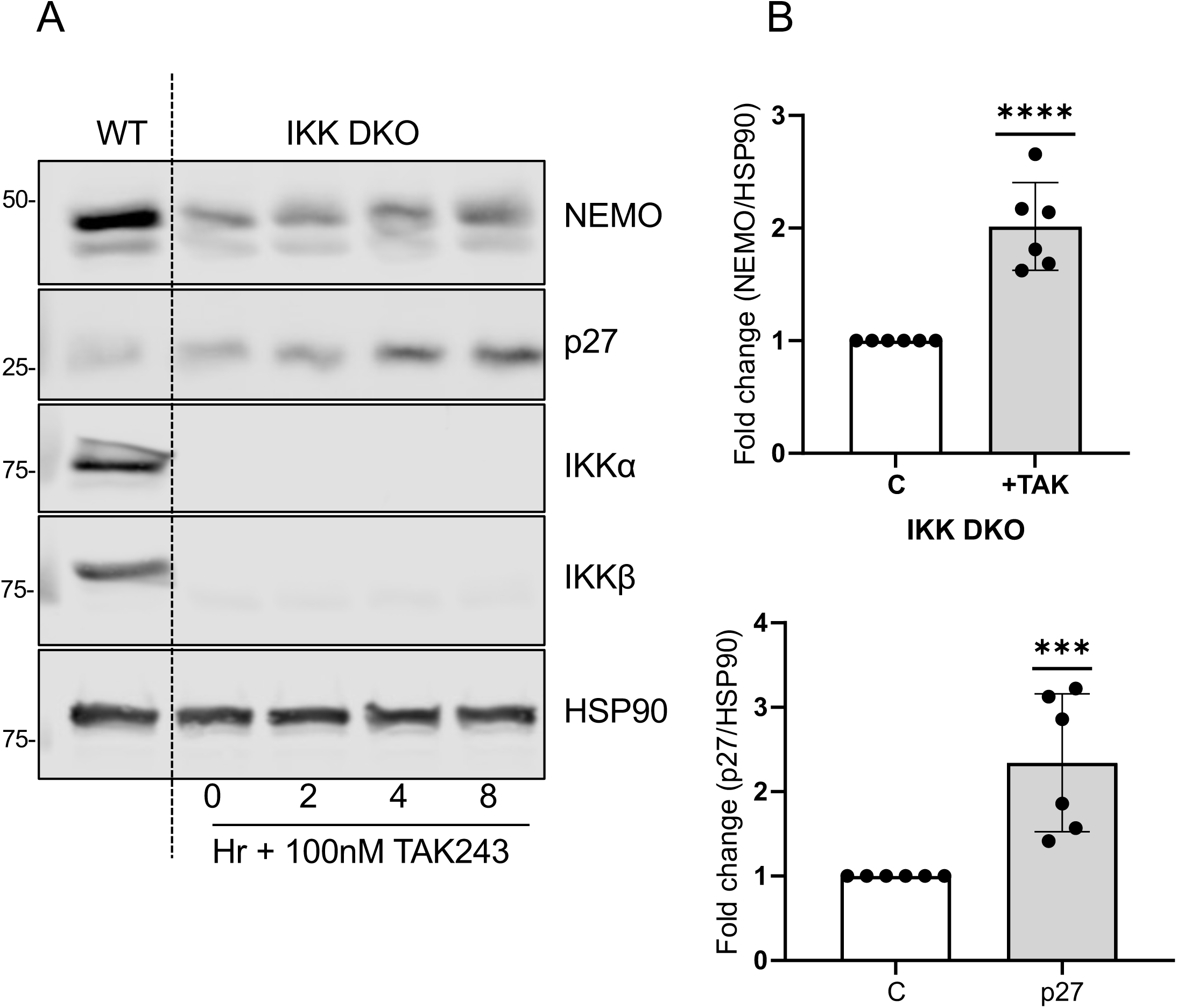
The UBA1 E1 ubiquitin-activating enzyme inhibitor TAK-243 increases the abundance of ENMO and p27^KIP1^ in IKK DKO cells. **(A)** IKKα/β DKO cells were exposed to 100nM TAK243 for the indicated times. Whole-cell lysates were fractionated by SDS-PAGE and Western blotted with the indicated antibodies. Data are from a single representative experiment. **(B)** Data for the untreated control or 8 hours TAK243 from n=6 experiments were quantified by LiCor and NEMO or p27 band intensity relative to HSP90 loading control calculated. Results are shown as mean ± SD. One sample t-test was carried out with Wilcoxon’s analysis (****=p<0.0001; ***=p<0.001).

### NEMO contains intrinsic disorder regions and can be degraded by 20S proteasomes

The minimal signal required for proteasome targeting is a chain of four K48-linked ubiquitin molecules, although other ubiquitin linkages can also target proteins for degradation [32,33]. However, ubiquitin modification of a folded protein alone is not sufficient for rapid proteasomal degradation; an intrinsically disordered region (IDR), either terminal or internal, to which the proteasome binds and initiates degradation, is also required [34]. Notably, many intrinsically disordered proteins (IDPs), such as Rb, p21^CIP1^, p53, c-Fos and BIM_EL_ can undergo ubiquitin/ATP-independent degradation by the 20S core proteasome both *in vitro* and *in vivo* [35–39]. Such proteins typically have IDRs towards their N- or C-termini, which can target the protein for degradation without the need for ATP-dependent unfolding [40].

Using a range of protein disorder prediction algorithms we identified major regions of intrinsic structural disorder within NEMO, with a striking level of agreement between the various predictors (Figure 7A-B and Supplementary Figure 4A-C). Our analysis indicated two long (> 30 aa) high confidence IDRs, one at the extreme N-terminus, and one near the C-terminus. A number of shorter, internal IDRs were also predicted, but with less agreement on their location. Seven of eleven prediction tools identified a high confidence 20aa IDR centred on aa 180, while six of eleven tools, predicted a high confidence 20aa IDR centred on aa 250. The structure/disorder consensus taken from the Protein data bank (PDB) structures available for NEMO indicated that the predicted IDRs were found approximately between the regions of solved structure (Figure 7A). This was also reflected in the plot of non-assigned X-ray structure coordinates, as defined by REMARK-465 entries in the PDB (Figure 7B). The difficulty in crystallising full-length NEMO protein may reflect these significant regions of intrinsic disorder. Prompted by the IDR predictions in NEMO we investigated whether NEMO might be degraded in a Ub-independent manner by 20S proteasomes. [^35^S]-methionine labelled NEMO was synthesised in an *in vitro* coupled transcription and translation (T&T) reaction and incubated with purified 20S proteasomes. NEMO was rapidly degraded within 60 minutes of incubation with 20S proteasomes (Figure 7C & 7D).

**Figure 7.**
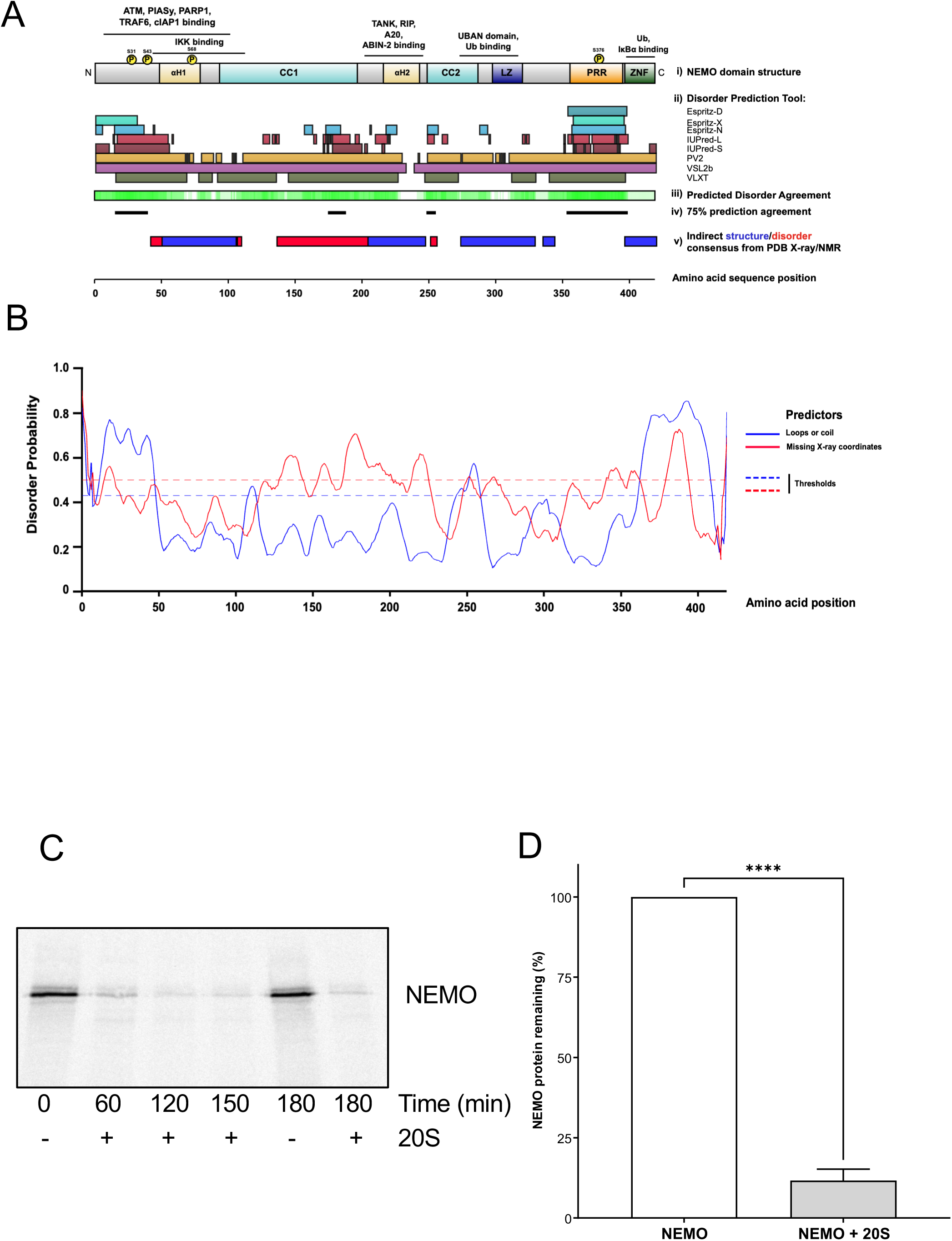
NEMO exhibits regions of intrinsic disorder and is degraded by 20S proteasomes *in vitro*. **(A) (i)** A diagram of key functional domains found in NEMO together with known/predicted protein interaction sites and identified (either in vitro or in vivo) IKKβ-specific phosphorylation sites. **(ii)** On the same amino acid sequence scale are the results of a selection of protein disorder predictors (Espritz, IUPRED, PV2, VSL2b, VLXT) taken from the D2P2 database. Coloured blocks represent disordered regions having a confidence score higher than 0.5, with a 5% false positive rate. **(iii)** The level of agreement between all of the disorder predictors is shown in green. **(iv)** Directly beneath this is an indication of sequence positions for which there is 75% agreement between all predictors in the database for the region being disordered. **(v)** Below this is a representation of consensus regions of defined structure or disorder provided by the various X-ray and NMR structures of NEMO within the PDB (consensus dataobtained from MobiDB database). **(B)** Intrinsic disorder probability plot for NEMO generated using DisEMBL. Loops/coils (in blue) as defined by DSSP -alpha-helix, 3-10 helix or beta-strand defined as ordered, and all other states as loops/coils. Missing X-ray coordinates (red), as defined by REMARK-465 entries in PDB. Dashed lines mark the threshold above which amino acids are regarded as disordered according to the two separate prediction definitions. **(C)** Flag-NEMO was synthesized *in vitro* using a T&T reaction in the presence of [^35^S]-methionine, followed by incubation with purified 20S proteasomes for the indicated time intervals. Samples were fractionated by SDS-PAGE and detected by autoradiography. **(D)** Results quantified from three independent experiments in which Flag-NEMO was incubated with 20S proteasomes for 2 hours. Statistical testing performed using unpaired, two-tailed Student’s t-test (*** =p<0.001).

Proteins with significant IDRs are thought to be inherently sensitive to proteasomal degradation but are often protected by their interaction with binding partners [41]. Such interactions typically occur through functional elements known as molecular recognition features (MoRFs), short amino acid sequences within longer disordered regions that undergo disorder-to-order transitions upon binding to their interaction partners [42]. We used a selection of the most accurate MoRF identification tools with differing prediction strategies to search for potential MoRFs within NEMO (Figure 8A-C; Supplementary Figure 4D & 4E). All prediction tools identified at least one high confidence MoRF within the N-terminal 50aa of NEMO; there was strong agreement on the presence of at least one high confidence MoRF within the C-terminal 50aa; and two predictors, ANCHOR and MoRFPred (Figure 8A & 8B), identified a high confidence MoRF within the leucine zipper (LZ) domain of NEMO. Interestingly, each of the prediction tools identified at least one potential MoRF within the minimal IKK binding domain (IBD) of NEMO (aa 44-111). The IKKs can protect NEMO from degradation in cells (Figure 1) so to investigate whether IKKs might protect NEMO from degradation by 20S proteasomes in vitro, NEMO was incubated with or without IKKα or IKKβ in the presence of purified 20S proteasomes; both IKKα and IKKβ significantly protected NEMO from 20S-dependent degradation (Figure 8D & 8E). The greater protection mediated by IKKβ compared to IKKα was at odds with the observation that IKKα KO cells exhibited a greater reduction in NEMO protein than IKKβ KO cells. However, the greater band smear observed when IKKα was co-expressed with NEMO, compared to IKKβ, may indicate that the IKKα was also partially degraded under these conditions.

**Figure 8.**
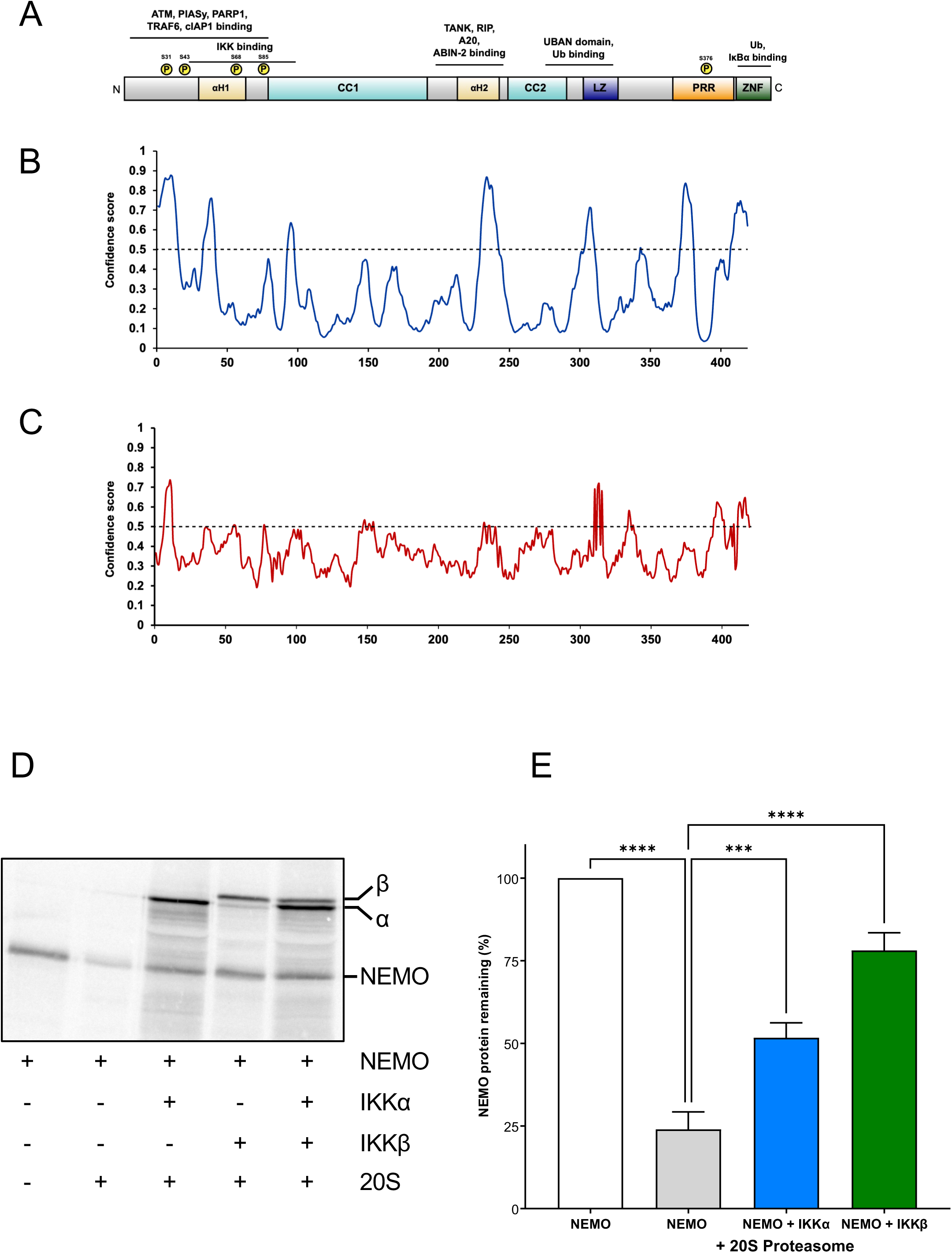
Inclusion of recombinant IKKα or IKKβ protects NEMO from degradation by 20S proteasomes. **(A)** A diagram of key functional domains found in NEMO together with a known/predicted protein interaction sites and identified (either in vitro or in vivo) IKKβ-specific phosphorylation sites. **(B)** On the same amino acid sequence scale a plot of confidence scores derived from ANCHOR predictor that residues within NEMO are within molecular recognition features (MoRFs). Residues with confidence scores above threshold = 0.5 are predicted to be within MoRFs. **(C)** Plot of confidence scores derived from MoRFpred that residues within NEMO are within MoRFs). Residues with confidence scores above threshold = 0.5 are predicted to be within MoRFs. **(D)** Flag-NEMO, Flag-IKKα and Flag-IKKβ were synthesized in vitro using a T&T reaction and [^35^S]methionine. FLAG-NEMO was then incubated with purified 20S proteasomes for 1 hour with or without FLAG-IKKα or FLAG-IKKβ. The degradation reaction was stopped, samples were subjected to SDS-PAGE and detected by autoradiography. **(E)** Results quantified from three independent experiments. Statistical testing performed using one-way ANOVA with Tukeys multiple comparisons test (***=p<0.001).

Together these results demonstrated that NEMO exhibits extensive regions of disorder, some of which may be modulated by protein binding. Furthermore, recombinant NEMO was rapidly degraded by 20S proteasomes and this degradation could be prevented by interactions with its IKK binding partners.

### Low NEMO expression in IKK DKO cells is enhanced by re-expression of wild type or kinase-dead IKKα but this requires IKK binding to NEMO

The preceding results indicated that the IKKs play a critical role in stabilising NEMO and in their absence NEMO undergoes proteasome-dependent degradation. Whilst our early experiments were in cells stimulated with TNFα (Figure 1A) NEMO degradation in IKK KO cells was apparent regardless of stimulation (see Figure 1B), questioning the need for IKK activity for stabilisation of NEMO. To address this we re-expressed wild type or catalytically inactive (kinase-dead) IKKα in IKKα/β DKO cells alone or together with Flag-NEMO. The low expression of endogenous NEMO in IKKα/β DKO cells was clearly increased when wild type IKKα or kinase-dead IKKα were transfected (Figure 9A & 9B). Similar results were observed for Flag-NEMO when it was co-expressed with wild type or kinase-dead IKKα (Figure 9A & 9C). Thus, NEMO was partially restored by re-expression of IKKα and this did not require its kinase activity.

**Figure 9.**
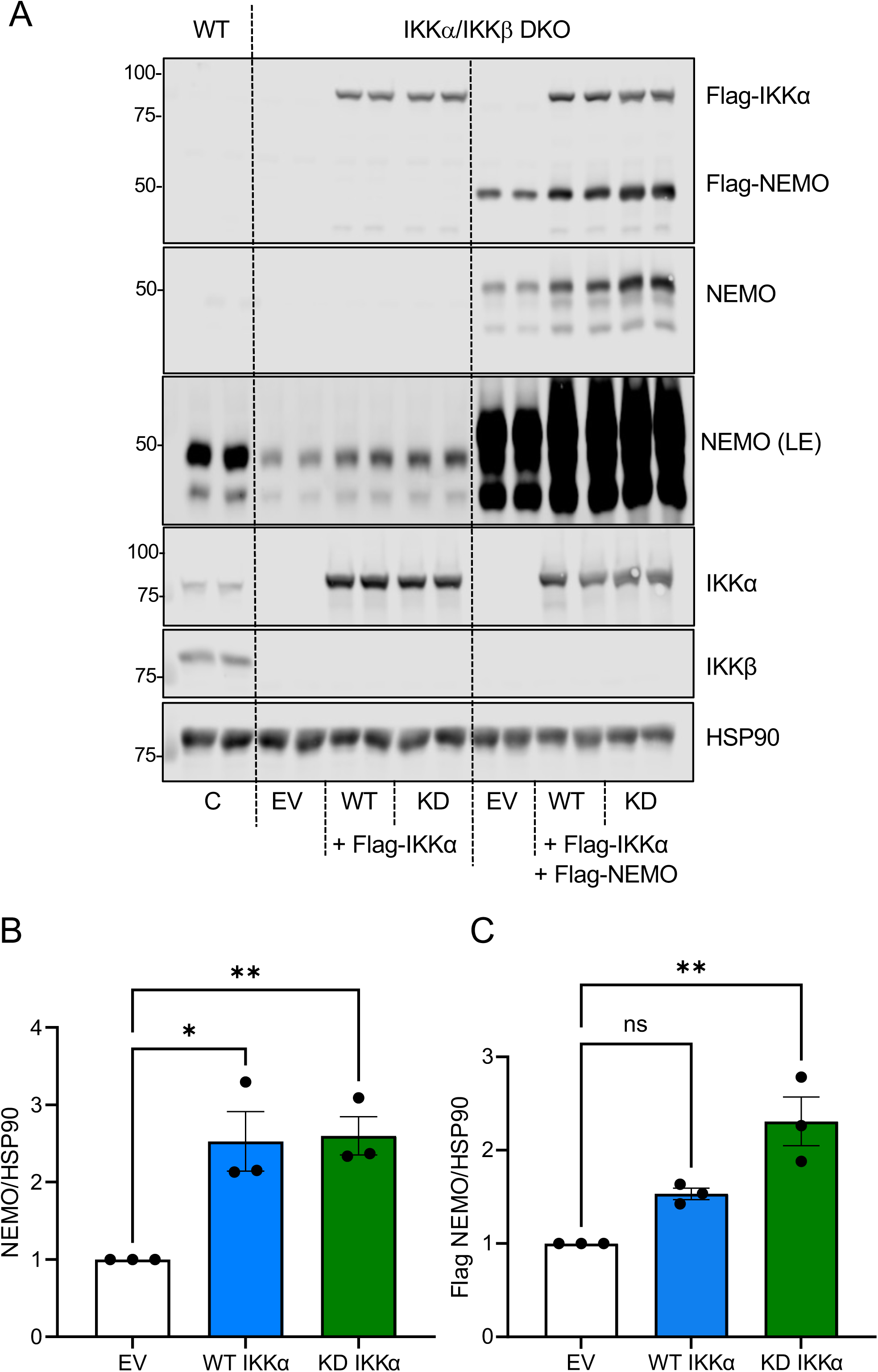
Low NEMO levels in IKK DKO cells are increased by re-expression of wild type or kinase-dead IKKα. **(A)** Duplicate dishes of IKKα/β DKO HCT116 were transfected with Flag-IKKα Wild type (WT) or Flag IKKα kinase dead (KD) with or without Flag-NEMO. Whole-cell lysates were fractioned by SDS-PAGE and Western blotted with the indicated antibodies. Data are from a single experiment representative of three giving identical results. **(B & C)** Data from n=3 experiments in (A) were quantified by LiCor imaging and endogenous NEMO (B) or Flag-NEMO (C) band intensity relative to HSP90 loading control calculated. Results are shown as mean ± SD of three independent experiments. One way ANOVA with Dunnett’s multiple comparison test was carried out between empty vector and either WT IKKα or KD IKKα (*=p<0.05; **=p<0.01). LE = Long Exposure.

Since IKK kinase activity was dispensable for stabilisation of NEMO, and IKKs can protect NEMO from degradation by 20S proteasomes in vitro, we reasoned that it was IKK binding to NEMO that was important. To test this we generated wild type or kinase-dead IKKα mutants lacking the C-terminal LDWSWL NEMO binding domain (ΔNBD) [15] and compared their NEMO-binding and NEMO-stabilising properties with WT IKKα. Co-immunoprecipitation experiments confirmed that the IKKαΔNBD mutants (wild type or kinase-dead) exhibited a substantial >90% reduction in binding to NEMO (Supplementary Figure 5). When the IKKαΔNBD mutants were expressed in IKK DKO cells they failed to increase the abundance of endogenous NEMO, and this applied for both wild type or kinase-dead IKKα (Figure 10A & 10B), indicating that direct binding of IKKs to NEMO is required to promote NEMO stability.

**Figure 10.**
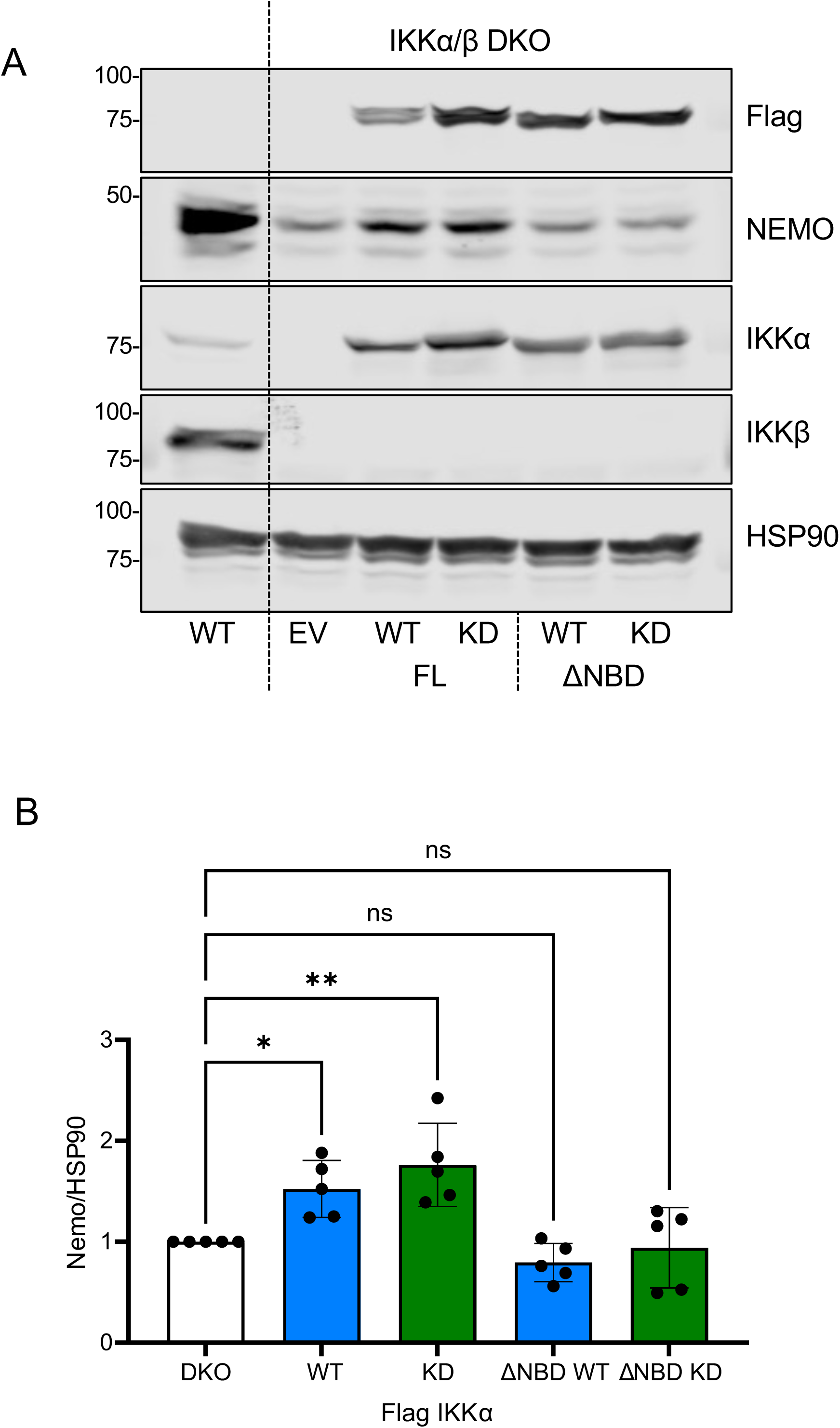
The NEMO binding domain (NBD) of IKKα is required to increase expression of NEMO in IKK DKO cells. **(A)** IKKα/β DKO HCT116 cells were transfected with either empty vector (Tag2b) or Flag-tagged full length or C-terminally truncated (ΔNBD) WT or kinase dead (KD) IKKα. Twenty-four hours post transfection whole cell lysates were prepared, fractionated by SDS-PAGE and Western blotted with the indicated antibodies. Data are from a single experiment representative of 5 giving similar results. **(B)** Data from n=5 experiments in A were quantified by LiCor imaging and NEMO band intensity relative to HSP90 loading control calculated. Results shown are mean ± SD. One way ANOVA was carried out with Dunnett’s multiple comparison test (*= p<0.05; **=p<0.01).

## DISCUSSION

Whilst NEMO is an essential scaffold for canonical IKK activation, our results suggest that the IKKs are in turn essential scaffolds that stabilise NEMO. In IKK DKO HCT116 cells the half-life of NEMO was greatly accelerated and steady-state NEMO protein abundance was 90% lower than WT cells. A similar reduction in NEMO was observed in IKKα KO SW620 cells and was enhanced with IKKβ siRNA. Studies of IKKα and IKKβ knockout mouse embryonic fibroblasts (MEFs) did not previously report loss of NEMO protein [43–45]. This discrepancy could reflect differences between MEFs and human cancer cell lines, but the conservation of NF-κB pathway between humans and mice makes this unlikely. A more prosaic explanation is that studies that performed knockout or knockdown of IKKα and/or IKKβ often neglected to probe for NEMO expression [46–49] or, when they did, NEMO western blots were over-exposed, inadvertently concealing a reduction in NEMO expression. Evidence to support our observations comes from a rare form of severe combined immunodeficiency (SCID) caused by a premature stop codon in *IKBKB*, encoding IKKβ, that results in loss of IKKβ protein [50]. Fibroblasts and peripheral-blood mononuclear cells from these patients exhibit a complete loss of IKKβ protein accompanied by a significant loss of IKKα and NEMO protein, providing confidence in the relevance of our results.

The reduction in NEMO expression in SCID patients with *IKBKB* mutation was not due to reduced mRNA expression and the underlying mechanism was not resolved [50]. Our results show that loss of IKKs drives the proteasome-dependent turnover of existing NEMO protein. Whilst lysosomal degradation of NEMO has been described [51–53], this is unlikely to account for the loss of NEMO described here, since Chloroquine disrupted lysosome function (shown by increased basal LC3B levels) but did not increase NEMO abundance in IKK DKO cells. Studies have suggested that NEMO may be degraded via the proteasome, but these studies were unrelated to IKK abundance and/or based on relatively modest evidence. A study showed that the HSP70 co-chaperone, BAG3, was required to maintain NEMO levels [54] though the authors used 200nM BAG3 siRNA, a concentration at which off-target effects are highly likely. HSP70 does interact with NEMO and inhibit its oligomerization *in vitro* [55,56], but HSP70-dependent delivery of NEMO to the proteasome has not been demonstrated. HSP90 is required to prevent proteasomal degradation of nascent IKKα and IKKβ [57–58], but is not required for NEMO stability [58], arguing against a critical role for HSP90 in control of NEMO abundance.

Whilst NEMO is degraded via the proteasome in IKK DKO cells, the identity of the substrate-specifiying E3 ubiquitin ligase remains unknown. SAG (Sensitive to apoptosis gene), also known as RBX2 or ROC2, a core component of the SCF E3 ubiquitin ligase complexes interacts with NEMO [59] but our results argue against a role for cullin-based E3 ligases since MLN-4924 had no effect on NEMO turnover. The Classical Swine Fever Virus NS5A protein also drives ubiquitylation and degradation of NEMO, by an unknown E3 ligase, to inhibit host immune signalling [60], whilst late phase bacterial or viral infection promotes an interaction between NEMO and MARCH2 (a RING-finger E3 ligase) driving K48-linked ubiquitylation and degradation of NEMO [61]. However, both scenarios are very different from the loss of the IKKs described here and siRNA-mediated knockdown of MARCH2 had no effect on NEMO abundance (data not shown).

In addition to ubiquitin-dependent degradation by 26S proteasomes our analysis identified intrinsically disordered regions (IDRs) in NEMO leading to the discovery that NEMO, like many other intrinsically disordered proteins (IDPs), can be degraded directly by 20S proteasomes in the absence of polyubiquitylation. The mechanism by which proteins are degraded by the 20S without ubiquitylation is poorly defined but may involve unstructured termini entering through the narrow gate of the 20S catalytic core during stochastic gate openings [62,63]. Sensitivity to 20S degradation has been proposed as an operational definition for IDPs [64] and it is notable that yeast, mouse and human proteins with terminal or internal IDRs have significantly shorter half-lives than proteins without such features [65]. Notably, re-expression of IKKα could partially restore NEMO expression in IKK DKO cells; this did not require IKK kinase activity but did require the NEMO binding domain of IKKα. Similarly, IKKα or IKKβ inhibited degradation of NEMO by 20S proteasomes *in vitro* suggesting that it is the interaction with IKKs that protects NEMO from degradation. These results are consistent with the ‘nanny’ model in which partner proteins protect IDPs from ‘degradation by default’ within the cell by masking or induce folding of IDRs that would otherwise target them for 20S-mediated degradation [66]. Our results suggest that NEMO contains significant regions of intrinsic disorder at its N- and C-termini that may act as effective initiation sites for degradation by 20S proteasomes in the absence of its binding partners, IKKα and IKKβ. This in turn raises the question of whether cells use 20S, 26S or both to degrade NEMO in the absence of the IKKs. The ability of MG132 to inhibit NEMO turnover in IKK DKO cells does not distinguish between these scenarios as it inhibits the catalytic core that is common to 20S and 26S. TAK-243, a highly selective inhibitor of the UBA1 E1 ubiquitin-activating enzyme, increased basal NEMO levels suggests that the canonical ubquitylation cascade does contribute to NEMO degradation in IKK DKO cells. However, in emetine chase experiments TAK-243 did not alter the rate of NEMO turnover.

Taken together our results suggest that NEMO may be degraded by both ubiquitylation-dependent (26S) and ubiquitylation-independent pathways in IKK DKO cells; the latter may involve uncapped 20S proteasomes or 20S proteasomes with alternate caps such as PSME1 (PA28α), PSME2 (PA28β), PSME3 (also known as PA28γ) or PSME4 (also known as PA200) that are selective for disordered proteins [67]. Such redundancy suggests a pressure to ensure NEMO degradation in the absence of the IKKs, but why? NEMO is a scaffold and a regulatory hub; its elongated structure facilitating simultaneous interaction with numerous protein partners and different forms of ubiquitin chain [14, 20], thereby bringing many proteins into the IKK signalosome; indeed, protein microarrays have identified over 100 NEMO interactors including several protein kinases that are not part of the core IKK signalosome [59]. Such promiscuity is a characteristic of IDPs and serves their common function as regulatory hubs within protein interaction networks [41, 65]. However, this promiscuity can be harmful if not regulated [68]; indeed, ∼80% of oncogenes contain predicted IDRs of 30 residues or longer [69], while an altered abundance of IDPs is commonly associated with several disease states, most notably α-synuclein and Tau proteins in Parkinson’s and Alzheimer’s disease [70–72]. ‘Degradation by default’ may ensure that the availability of proteins with significant IDRs that have become uncoupled from their normal functional complexes is tightly controlled [73]. In this context, the rapid turnover of ‘IKK-free’ NEMO may prevent any undesirable interactions. It may also contribute to fine-tuning of NEMO protein stoichiometry in relation to the IKK subunits, such that downregulation of IKK expression is matched by enhanced degradation of NEMO protein. This model would dictate that functional interactions with NEMO would necessitate protection of NEMO from basal degradation. It is intriguing, therefore, that numerous proteins, including ATM, PIASy and PARP1, have been shown to interact with the N-terminal disordered region of NEMO in a manner mutually exclusive with IKKs.

Finally, our results have two further implications. First, we recently showed that loss of IKKα had a much greater effect on canonical NF-kB signalling than loss of IKKβ in colorectal cancer cells [23]. Whilst those results were unarguable, their interpretation could be clouded by the fact that loss of IKKα reduces NEMO abundance by 50-60%, which might have consequences for the remaining IKKβ-dependent NF-κB signalling. However, IKKβ S177/181 phosphorylation was actually enhanced in IKKα KO cells over both short and long time courses [23]; this is consistent with binding between NEMO and IKKα being weaker than with IKKβ [15,18], so that loss of IKKα facilitates IKKβ binding to NEMO and IKKβ activation. This in turn suggests that even when NEMO is reduced by 50-60% there is still sufficient to support IKKβ activation. Despite this adaptive enhancement of IKKβ activation, IKKα KO cells still exhibit reduced p65 NF-kB phosphorylation and nuclear translocation [23] suggesting p65 exhibits a dependency upon IKKα that cannot be fulfilled by IKKβ. Regardless, since loss of IKKα facilitates IKKβ binding to NEMO and IKKβ activation, why is this not sufficient to also protect NEMO from degradation in IKKα KO cells? Maintenance of IKKβ signalling in IKKα KO cells suggests there is enough IKKβ to bind NEMO and support IKKβ catalysis but not enough total IKK (α+β) to stabilise the total NEMO pool, which is a non-catalytic, stoichiometric function of IKKs. Such considerations are dependent on the absolute abundance of IKKα, IKKβ and NEMO and stoichiometry of IKK:NEMO complexes which clearly differ considerably between cell types [74]. Second, our results may have implications for therapeutic approaches to targeting the IKKs. IKKβ kinase inhibitors have failed to progress to clinical approval [75] and IKKα inhibitors are at an earlier stage [76]. Cell-penetrant peptides derived from the NBD of IKKβ have shown efficacy in models of inflammation but are not being progressed due to the expense of peptide synthesis and poor oral bioavailability [75]. Structure-guided screening has identified small molecule NBD-mimetics that show efficacy in inflammation and ageing-related disease models [77, 78]. These NBD mimetics disrupted NEMO:IKKβ interactions but had little effect on NEMO:IKKα interactions and did not lower NEMO abundance; however, this was based on a 30 minute cell treatment which is likely too short for significant NEMO turnover. Will longer term treatment with IKKβ-derived NBD mimetics drive NEMO loss or will it drive a greater dependence on the remaining NEMO:IKKα which might serve as a resistance mechanism or a vulnerability?

In summary, loss of IKKα and IKKβ drives the proteasomal degradation of NEMO via ubiquitylation-dependent and independent pathways, the latter consistent with the presence of regions of disorder at the N-terminus of NEMO which overlap with the IKK binding site. Our results indicate that as well as their catalytic function, IKKα and IKKβ act as critical kinase-independent scaffolds that stabilise NEMO by direct binding. Since NEMO serves as a signalling hub, interacting with >100 proteins including several other protein kinases that are not part of the core IKK signalosome [59] these results may also have wider implications beyond the NF-κB pathway.

## MATERIALS AND METHODS

### Cell Culture and Constructs

HCT116 cells were provided by Bert Vogelstein and cultured in high glucose DMEM supplemented with 10% (v/v) FBS, penicillin (100 U/mL), streptomycin (100 mg/mL), and 2mmol/L glutamine (all reagents Life Technologies). IKKα KO SW620 cells were derived as previously described (Prescott et al 2022) and were cultured in Leibovitz L15 medium supplemented with 10% (v/v) FBS, penicillin (100 U/mL), streptomycin (100 mg/mL), 2 mmol/L glutamine and 0.075% sodium bicarbonate (v/v). Cells were maintained in a humidified incubator at 37°C 5% CO_2_ (v/v). Flag tagged IKKα WT and KD were a gift from Prof Lienhard Schmitz (Department of Biochemistry, Justus-Liebig-Universität Gießen, Germany. pCMV2TAG IKKβ WT and KD (K44M) were a gift from Anjana Rao (Addgene plasmid #11103 and #11104 respectively). pCMVTAG2B NEMO was a gift from Jon Ashwell (Addgene plasmid # 11970). pcDNA3 HA human NEMO was a gift from Kunliang Guan (Addgene #13512). Transfection of plasmids was carried out using JetPrime (#10100015 Sartorius) following the manufacturer’s instructions. C-terminal truncation of WT and KD IKKα was carried out using New England Biolabs (NEB) Q5 mutagenesis kit (E0554S) and NEB base changer software was used to design the required oligos following the manufacturer’s instructions. Oligo sequences are available on request.

### siRNA

Cells were transfected with 20nM siRNA smart pool as indicated (IKKα (CHUK) siGENOME smart pool M-003473-02, IKKβ (IKBKB) siGENOME smart pool M003503-03, NEMO siGENOME smart pool M-003767-02) or NT siRNA (siGENOME non targeting pool D-001206-13-20) using Lipofecatmine RNAiMAX transfection reagent, following the manufacturer’s protocol (Thermo Fisher #13778100). Forty-eight or 72 hours post transfection cells were lysed in TG lysis buffer and lysates prepared for SDS-PAGE and western blotting.

### NF-kB Luciferase reporter assays

Cells were seeded in triplicate in antibiotic-free media in 96-well white-walled plates (Thermo). The following day cells were transiently transfected with 100 ng pGL4.32 luc2P/NF-κB-RE luciferase reporter plasmid and 10 ng constitutively expressed pCMV renilla luciferase plasmid (used for normalisation) per well. 24 hours after transfection, treatments were performed as described. Luciferase activity was measured using the Dual-Luciferase Reporter Assay System (Promega), and on a FLUOstar Omega plate reader (BMG Labtech), according to the manufacturer’s protocol.

### NEMO CRISPR Gene editing

The three CRISPR guides for NEMO knockout were designed using Horizon’s discovery design tool and were 5’-GCAGCAGATCAGGACGTACT-3’, 5’-AGGTGCCTATTCATCCAACA-3’ and 5’-CTCCTCCAGGCAGCGCTGGA-3’. Guides were cloned into pSpCas9(BB)-2A-GFP (PX458) (a gift from Feng Zhang Addgene #48138) following the Ran *et al.* protocol [79] and the resulting constructs sequenced by Sanger sequencing to confirm correct insert. HCT116 cells were transfected with 5µg of each guide RNA construct using Jetprime transfection reagent following the manufacturer’s instructions. 48 hours after transfection GFP-positive cells were sorted by flow with one cell per well seeded onto 96 well plates and surviving clones grown up. NEMO knockout was confirmed by western blot.

### RT-qPCR

Total RNA was extracted from cells using the RNeasy Kit (QIAGEN #74104), QIAshredders (QIAGEN #79654) and RNase-Free DNase Set (QIAGEN #79254) according to the manufacturer’s instructions. 1 μg RNA was reverse transcribed to cDNA using the Quantitect Reverse Transcription kit (QIAGEN #205311) according to manufacturer’s instructions. cDNA was diluted 1:20 in RNase-free water and stored at -20°C. RT-qPCR was performed using SYBR green universal master mix (InVitrogen #4309155) and primers with the following sequences: NEMO FWD: 5’-GACAACCACATCAAGAGCAGC-3’ REV: 5’-CTTGTAGATATCCGCCTGGG-3’. YWHAZ FWD: 5’-TGATCCCCAATGCTTCACAAG-3’ REV: 5’-GCCAAGTAACGGTAGTAATCTCC-3’. UBC FWD: 5’-CTGGAAGATGGTCGTACCCTG-3’ REV: 5’-GGTCTTGCCAGTGAGTGTCT-3’. This was performed in a 96-well plate format using a CFX96 Real-Time PCR Detection System (BioRad). Data were normalised to the geometric mean of the reference gene expression and relative gene expression change calculated using the Pfaffl method. Statistical analysis was performed with the REST 2009 software package using an applied two-sided pair-wise fixed reallocation randomisation test.

### SDS-PAGE and Western blotting

When required cells were lysed in ice-cold TG-lysis buffer, centrifuged to remove insoluble material and assayed for protein content (Bradford). The normalised lysates were fractionated by SDS-PAGE as previously described followed by Western blotting with HRP-conjugated secondary antibodies and ECL detection was performed as described previously [80]. To enable quantification of western blots fluorescently tagged secondary antibodies and a Li-Cor Odyssey CLx Imaging System (LI-COR Biosciences) were used. Briefly, membranes were cut to allow probing for multiple molecular weight proteins. Where necessary, multiple independent blots were performed using the lysate from each experiment. Quantification of the proteins of interest was carried out after normalization to a loading control (HSP90 or actin) using Image Studio 6 or later software. For some experiments quantification was carried out using ImageJ where blots were detected using ECL.

### HA immunoprecipitation

Cells were lysed in TG lysis buffer, cleared by centrifugation and protein content assayed and lysate either used for the ip or boiled in 4x sample buffer for input blots. Anti-HA magnetic beads (Pierce #88837) were prepared following the manufacturer’s instructions before addition of equal amounts of protein added and volume of lysate adjusted to be equal for all samples. Samples were then incubated at room temperature with end over end mixing for 30 minutes. The lysate was then aspirated, and beads washed 4 times in TG lysis buffer. Beads were boiled in 1x sample buffer before fractionation by SDS-PAGE and western blotting.

### Antibodies

NEMO Cell Signalling Technology (CST) #2695 and Abcam #ab178872, IKKα N-term antibody CST #2682 and C-term antibody Abcam #ab109749, IKKβ C-term antibody CST #2684 and Abcam N-term antibody #ab124957, p-IKKα/β (S176/180) CST #2697, IKBα CST #9242, p-IKBα (S32/36) CST #9246, p65 CST #8242, p-p65 (S486) CST #3039, p-p65 S536) CST #3033, p27 CST #3686, Actin #A5441 Sigma, HSP90 BD Transduction Labs #610418, LC3B CST #2775, FLAG-M2 #3165 Sigma, HA CST #3724. Secondary antibodies used were CST Dylight anti-rabbit 800 #5151S and anti-mouse 800 #5257S. The HRP conjugated secondary antibodies were from BioRad: Goat anti-mouse HRP #170-6516 and Goat anti-rabbit HRP #170-6515. ECL reagents were from Cytiva #RPN2209.

### *In vitro* 20S proteasomal degradation assay

*In vitro* transcription and translation was performed in the presence of [^35^S]-methionine using the TnT Quick Coupled Transcription/Translation System (Promega), according to manufacturer’s instructions. Plasmids employed were pCDNA3.1-NFLAG NEMO, pcDNA3.1-NFLAG IKKα, pcDNA3.1-CFLAG IKKβ. Purified 20S proteasomes were generated based on a method described by Beyette et al [81]. [^35^S]-methionine-labelled proteins translated *in vitro* were incubated with 0.35 or 1 μg of purified 20S proteasomes in 150 mM NaCl, 50 mM Tris HCl pH 7.5 and 1 mM DTT at 37°C for a defined time period. Reactions were then resolved by SDS-PAGE and visualized by autoradiography.

### Cell Treatments

MG132 (#C221) Chloroquine (#PHR1258) and Emetine (#E2375) were all purchased from Sigma. MLN4924 (#S7109) and TAK243 (#S8341) were purchased from Selleckchem. TNFα (210-TA) was purchased from R&D Systems. Where required cells were treated as indicated with inhibitor or vehicle before lysis and further processing for downstream analysis. Results are presented as the mean ± SD unless otherwise stated. Statistical analysis was carried out as indicated in the figure legends using GraphPad Prism 10 (Boston MA).

## DATA AVAILABILITY

All relevant data are contained within the main article and its Supplementary Files.

## COMPETING INTERESTS

Jack Prescott’s PhD was supported in part by AstraZeneca. Eleanor Griffith’s PhD was supported in part by Mission Therapeutics. All other authors declare no competing interest associated with the manuscript.

## FUNDING

Jack Prescott’s PhD was funded by a Biotechnology and Biological Sciences Research Council (BBSRC) CASE PhD studentship in collaboration with AstraZeneca. Eleanor Griffith’s PhD was a Biotechnology and Biological Sciences Research Council (BBSRC) CTP CASE PhD studentship in collaboration with Mission Therapeutics. Work in the Cook laboratory was supported by the BBSRC Project Grant to Simon Cook (BB/S017062/1) and Institute Strategic Programme Grants BBS/E/B/000C0433, BBS/E/B/000C0531, BBS/E/B/000C0533, BBS/E/B/000C0534, BBS/E/B/000C0535, from the Biotechnology and Biological Sciences Research Council which also supported KB and SJC.

## CRediT AUTHOR CONTRIBUTION

**Jack A. Prescott:** Conceptualization, Formal analysis, Validation, Investigation, Methodology, Writing — original draft. **Kathryn Balmanno:** Formal analysis, Validation, Investigation, Methodology, Writing — review and editing. **Nadav Myers:** Formal analysis, Validation, Investigation, Methodology. **Eleanor J. Griffiths:** Formal analysis, Investigation, Validation. **Yosef Shaul:** Formal analysis, Supervision Investigation, Writing — review and editing. **Simon J. Cook:** Conceptualization, Supervision, Funding acquisition, Writing — original draft, Writing — review and editing and Project administration,.

## ACKNOWLEDGEMENTS

This project was started by Jack Prescott, our friend and former PhD student, and this paper is dedicated to his memory. We thank colleagues in the Signalling Programme at the Babraham Institute and Prof Neil Perkins for useful comments and criticisms on this study and members of the Cook laboratory for their support and encouragement. We would also like to acknowledge the excellent advice and support of the Babraham Institute Flow Cytometry Facility.

## ABBREVIATIONS

**Supplementary Figure 1.**
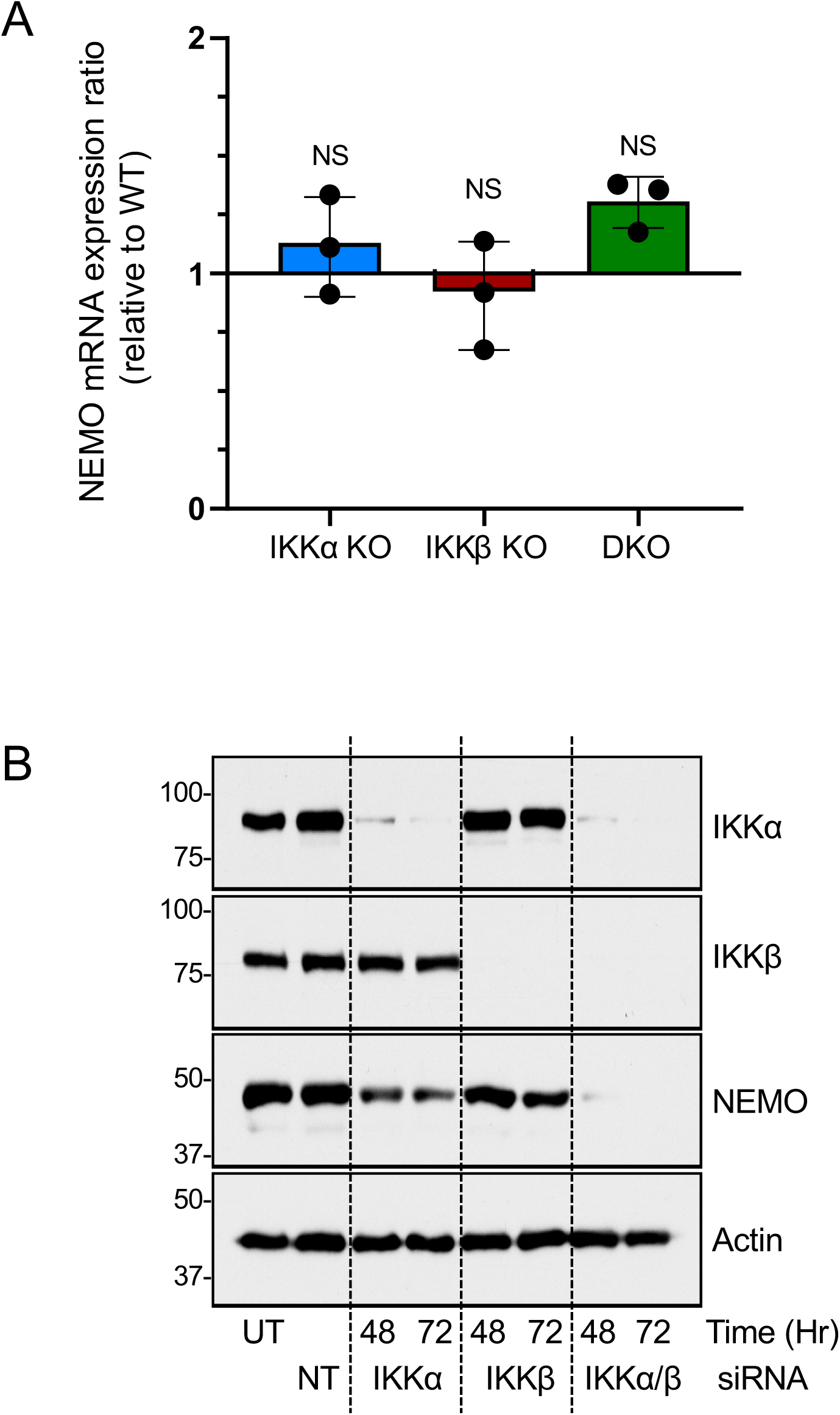
siRNA-mediated knockdown of IKKs reduces NEMO protein levels. **(A)** Three independently derived WT HCT116 clones and three clones each of IKKα single, IKKβ single and IKKα/β double knockout were seeded in normal growth medium for 48 hours prior to RNA extraction. Relative NEMO mRNA abundance was determined by RT-qPCR, with normalisation to the geometric mean of the reference gene (YWHAZ and UBC) expression. Data are plotted as mean ± SD relative to the WT control (line at 1 on the y-axis). One way ANOVA between WT and KO samples with Dunnett’s multiple comparison test revealed no significant difference (NS) in NEMO mRNA levels. **(B)** HCT116 cells were left untransfected (UT), transfected with non-targeting (NT) siRNA or transfected with either 20 nM IKKα-specific siRNA, 20 nM IKKβ-specific siRNA or 20 nM of both IKKα- and IKKβ specific siRNAs. Total siRNA was equalized to 40 nM with NT siRNA as required. 48 or 72 hours after transfection, whole cell lysates were fractionated by SDS-PAGE and Western blotted with the indicated antibodies. Data are from a single experiment with similar results obtained in two further experiments.

**Supplementary Figure 2.**
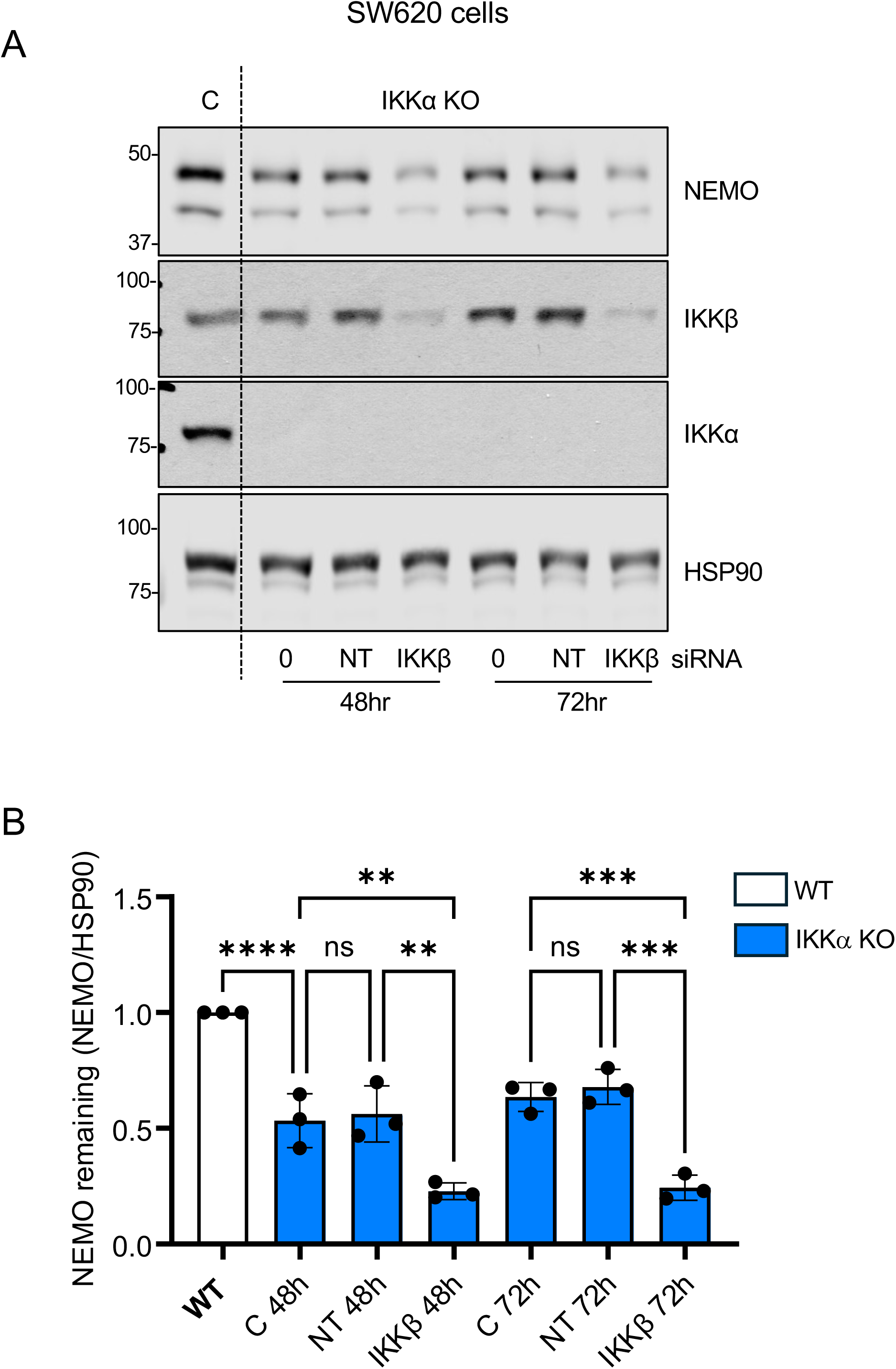
IKKα KO SW620 cells exhibit reduced NEMO levels that are further reduced by IKKβ knockdown. **(A)** IKKα KO SW620 cells were mock transfected or transfected with 20nM non-targeting (NT) or 20 nM IKKβ-specific siRNA siRNA for the indicated times. Whole cell lysates were fractionated by SDS-PAGE and Western blotted with the indicated antibodies. Results are from a single experiment representative of three giving similar results. **(B)** Data from n=3 experiments in (A) were quantified by LiCor imaging and NEMO band intensity relative to HSP90 loading control calculated, the mean ± SD are shown. One way analysis of variance with Tukey’s multiple comparison test was carried out (**=p<0.01; ***=p<0.001; ****=p<0.0001).

**Supplementary Figure 3.**
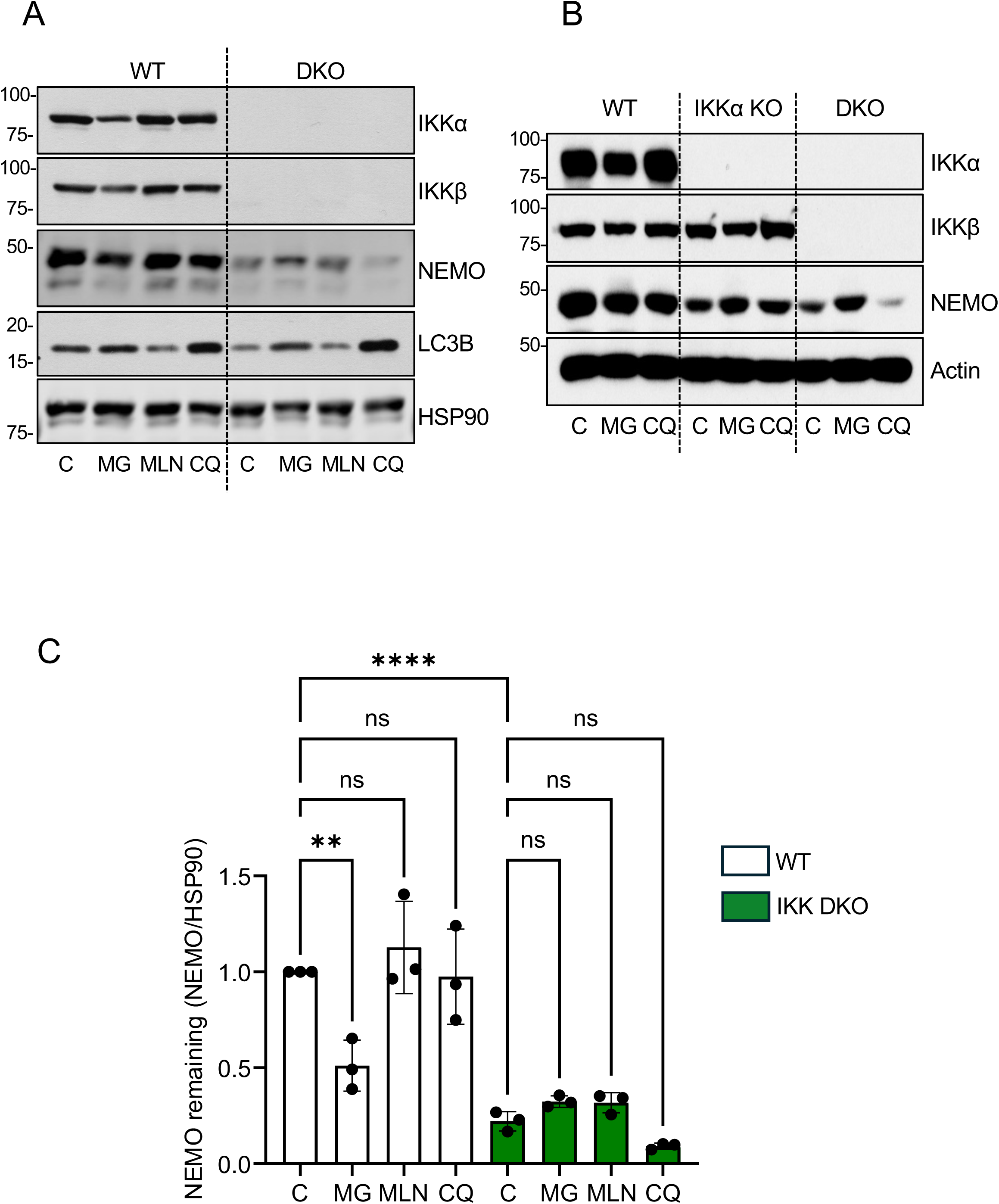
The effect of MG132, MLN4924 or Chloroquine on basal NEMO levels in WT, IKKα KO or IKK DKO cells. **(A)** IKK WT or IKKα/β DKO HCT116 cells were exposed to 7.5μM MG132 (MG), 300nM MLN (MLN4924) or 25μM Chloroquine (CQ) for 16 hours. Whole cell lysates were then fractionated by SDS-PAGE and Western blotted with the indicated antibodies. Results are from a representative experiment of three giving similar results. **(B)** IKK WT, IKKα KO or IKKα/β DKO HCT116 cells were exposed to MG132 or Chloroquine as in (A) and whole cell lysates fractionated by SDS-PAGE with Western blot analysis with the indicated antibodies. Results are from a representative experiment of 3 giving similar results. **(C)** Data from n=3 experiments in A were quantified by LiCor imaging and NEMO band intensity relative to HSP90 loading control calculated. Results shown are the mean ± SD. One way ANOVA with Tukey’s multiple comparison test was carried out (**=p<0.01; ****=p<0.0001).

**Supplementary Figure 4.**
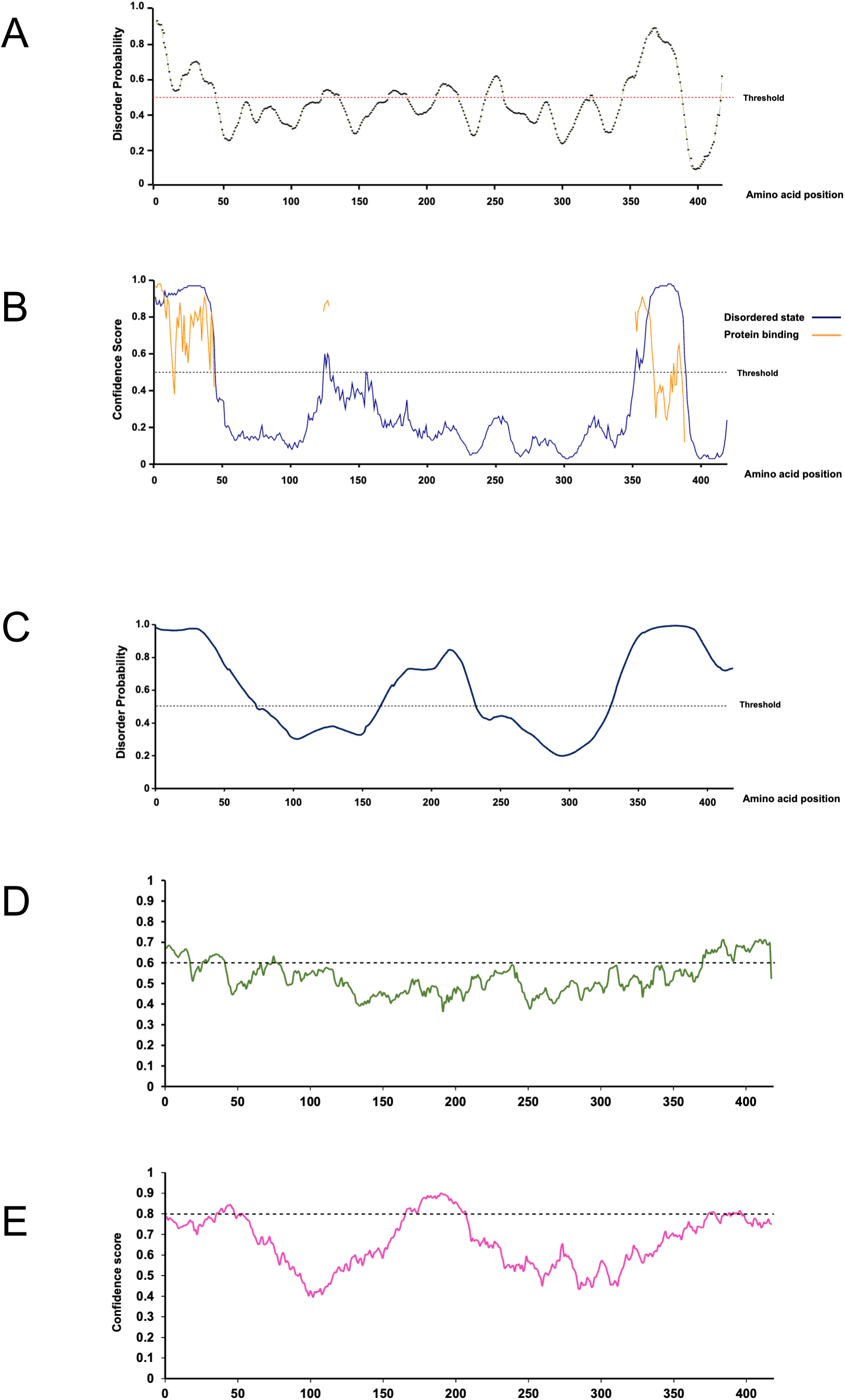
Bioinformatic analysis of disordered regions and MoRFs in NEMO. **(A)** Intrinsic disorder probability plot for NEMO generated using PrDOS. Threshold with 5% false positive rate shown as red dashed line. **(B)** Intrinsic disorder probability plot for NEMO generated using DISOPRED3. Blue line indicates residue disorder confidence levels. Threshold with 5% false positive rate shown as black dashed line. For disordered residues, the orange line shows the confidence of disordered residues being involved in protein-protein interactions. **(C)** Intrinsic disorder probability plot generated using MFDp2. Threshold with 5% false positive rate shown as black dashed line. **(D)** Plot of confidence scores derived from MoRFCHIBI SYSTEM predictor that residues within NEMO are within molecular recognition features (MoRFs). Residues with confidence scores above threshold = 0.6 are predicted to be within MoRFs. **(E)** Plot of confidence scores derived from DisoRDPbind predictor that residues within NEMO are within MoRFs. Residues with confidence scores above threshold = 0.8 are predicted to be within mORFs. In all cases **(A-E)** the same amino acid sequence scale x-axis represents the same NEMO same amino acid sequence scale.

**Supplementary Figure 5.**
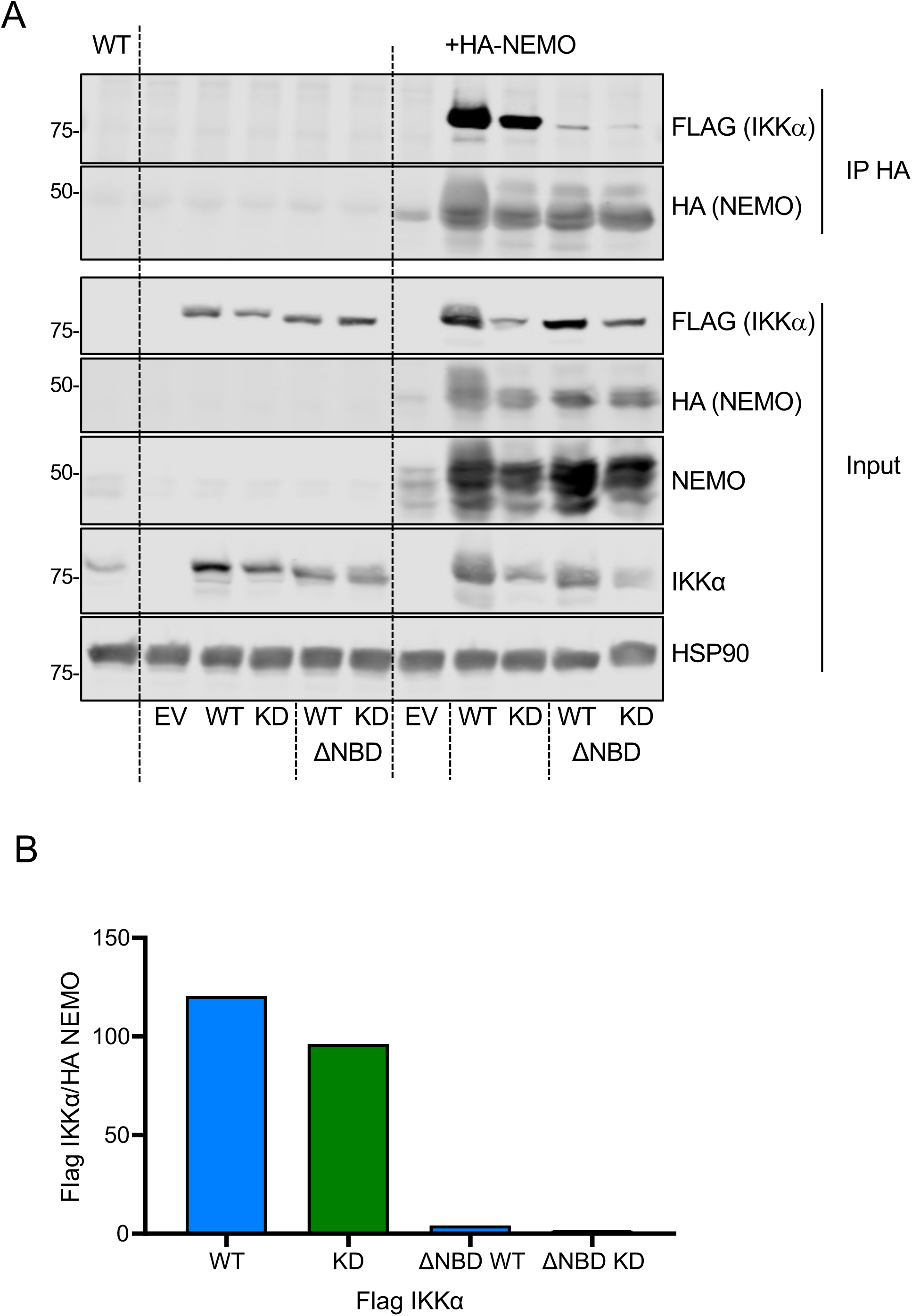
Confirmation that IKKα ΔNBD mutants are defective for NEMO binding. **(A)** IKKα/β DKO HCT116 cells were transfected with either empty vector (Tag2b) or Flag-tagged full length or C-terminally truncated (ΔNBD) WT or kinase dead (KD) IKKα with or without HA-tagged NEMO. Twenty-four hours post transfection cells were lysed and whole cell lysates subjected to immunoprecipitation using HA magnetic beads, with some lysate retained for input blots. Samples were fractionated by SDS-PAGE and Western blotted with the indicated antibodies. **(B)** Data from (A) was quantified by LiCor imaging and Flag (IKKα) band intensity relative to HA (NEMO) band intensity for the IP plotted.

## Notes

### Competing Interest Statement

The authors have declared no competing interest.

