## Supplementary Figures and legends for "IKKα and IKKβ serve as kinase-independent scaffolds that prevent proteasomal degradation of NEMO"

**Supplementary Figure 5. Confirmation that IKKα ΔNBD mutants are defective for NEMO binding. (A)** IKKα/β DKO HCT116 cells were transfected with either empty vector (Tag2b) or Flag-tagged full length or C-terminally truncated (ΔNBD) WT or kinase dead (KD) IKKα with or without HA-tagged NEMO. Twenty-four hours post transfection cells were lysed and whole cell lysates subjected to immunoprecipitation using HA magnetic beads, with some lysate retained for input blots. Samples were fractionated by SDS-PAGE and Western blotted with the indicated antibodies. **(B)** Data from (A) was quantified by LiCor imaging and Flag (IKKα) band intensity relative to HA (NEMO) band intensity for the IP plotted.

ity for the IP plotted.

A

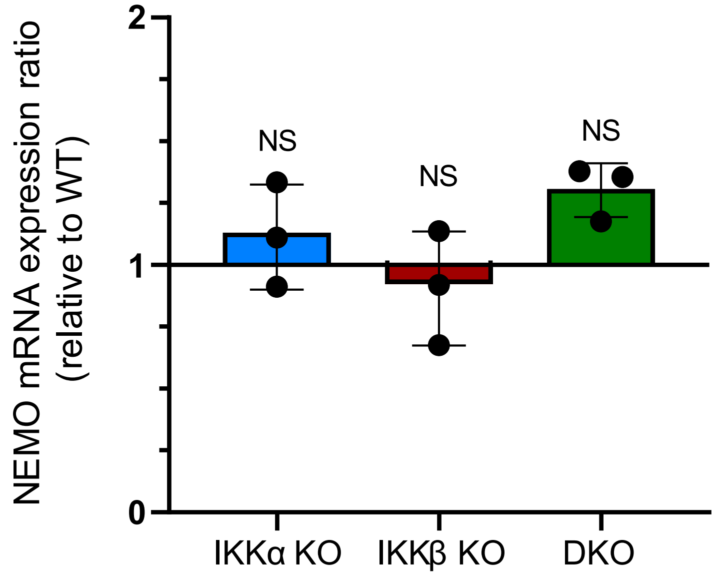
NEMO mRNA expression ratio (relative to WT)
2
1
0
NS
NS
NS
IKKα KO
IKKβ KO
DKO

B

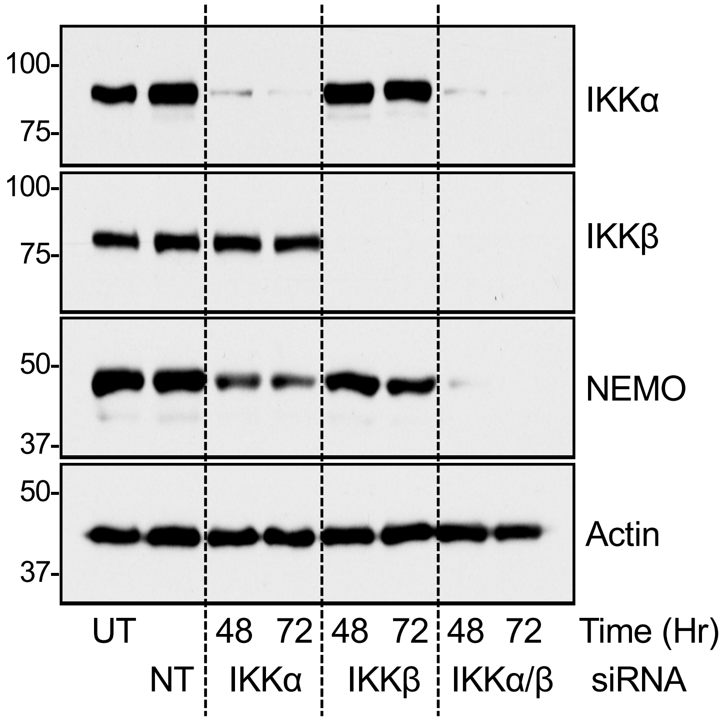
100-
75-
100-
75-
50-
37-
50-
37-
IKKα
IKKβ
NEMO
Actin
UT
48
72
48
72
48
72
Time (Hr)
NT
IKKα
IKKβ
IKKα/β
siRNA

Supplementary Figure 2

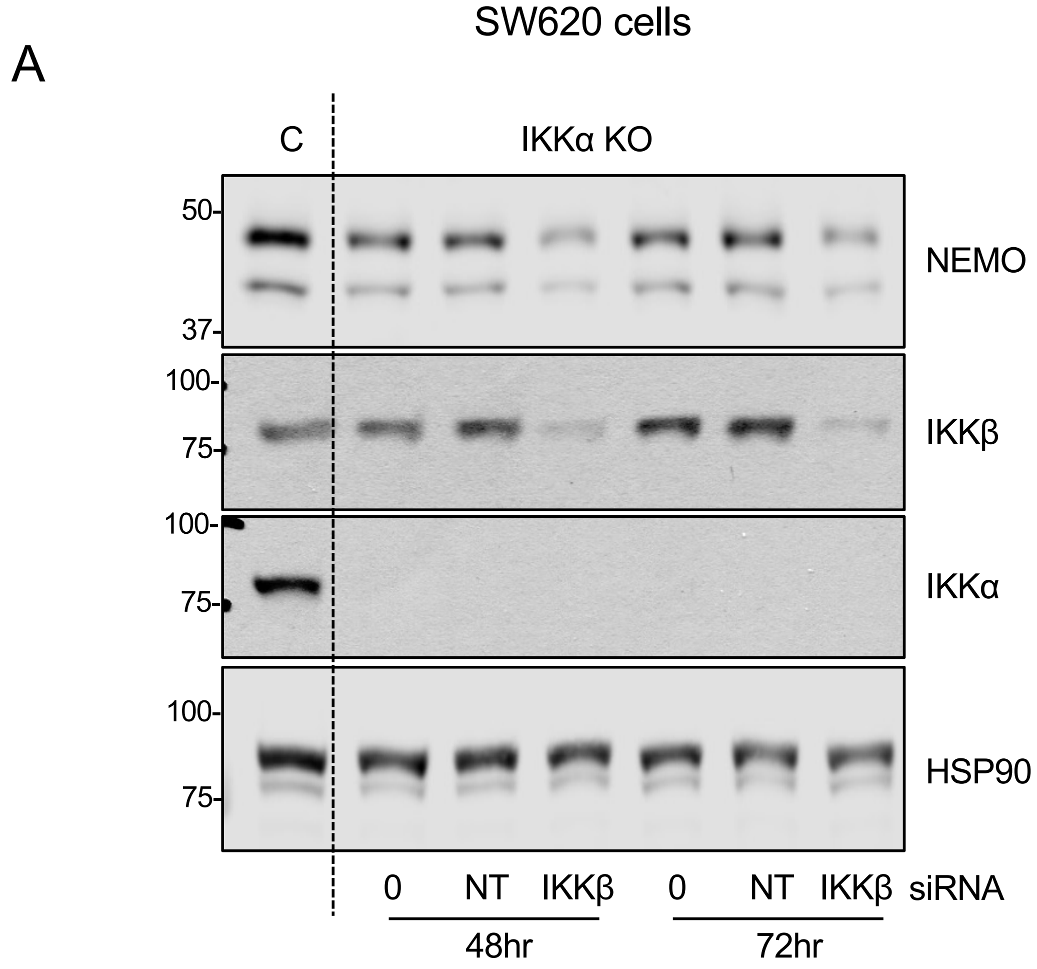


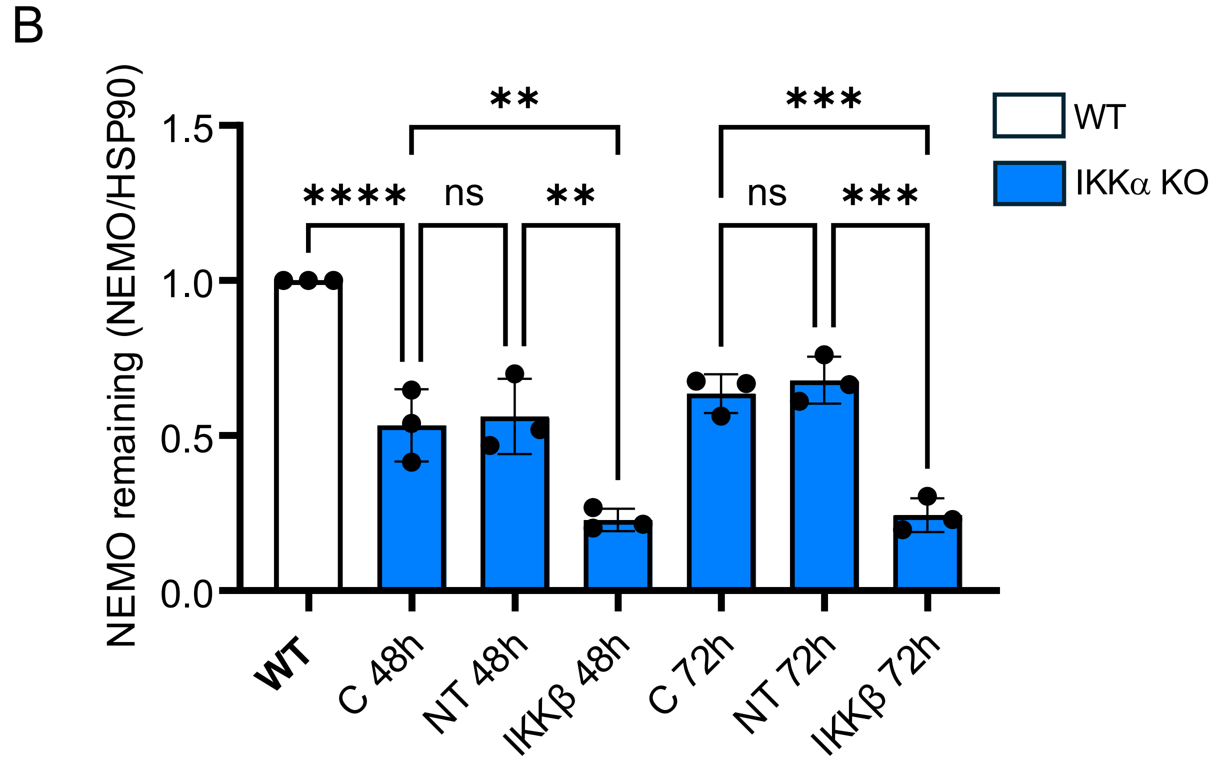

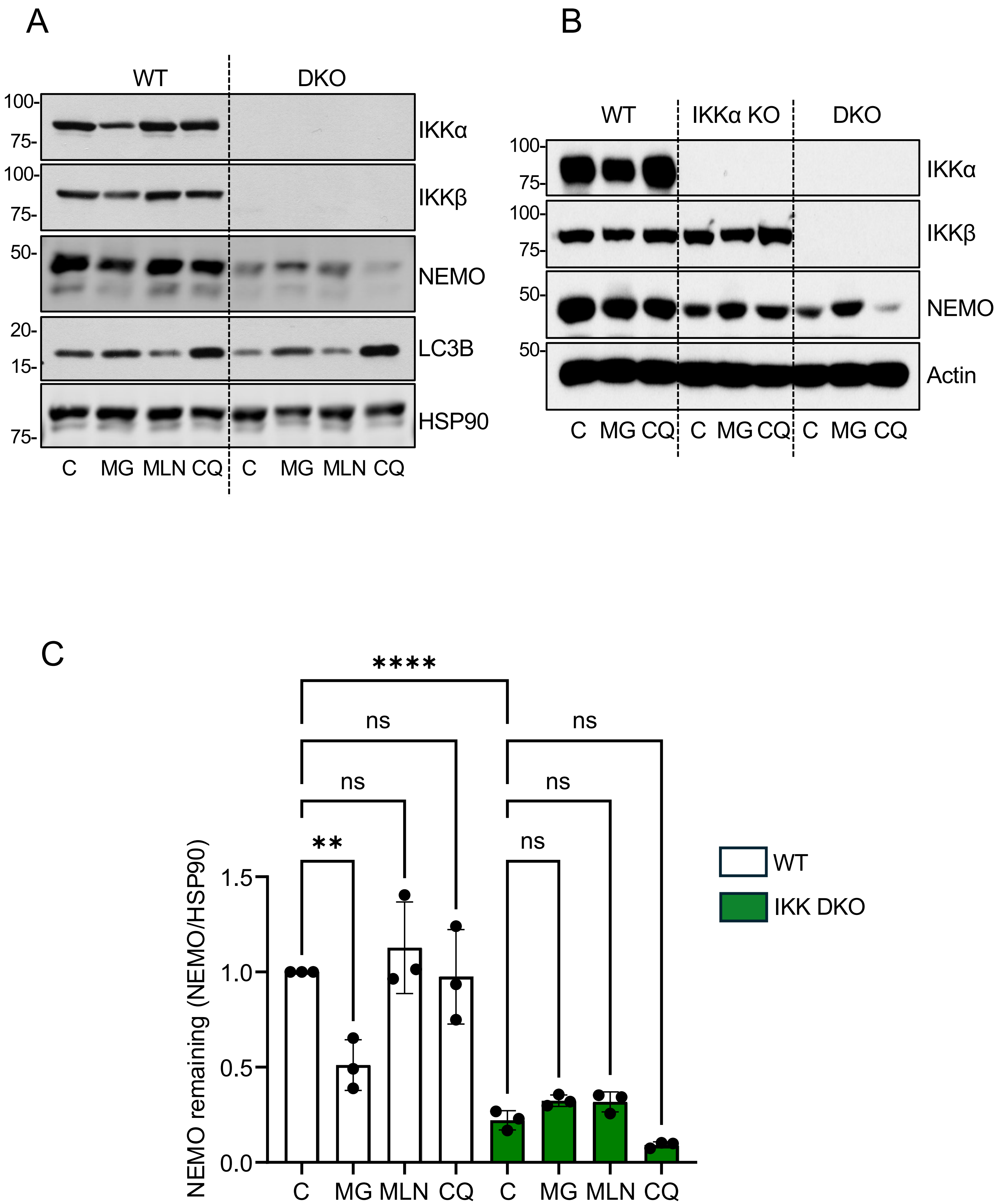
A
WT
DKO
100-
75-
IKKα
100-
75-
IKKβ
50-
NEMO
20-
15-
LC3B
75-
HSP90
C MG MLN CQ C MG MLN CQ
B
WT
IKKα KO
DKO
100-
75-
IKKα
100-
75-
IKKβ
50-
NEMO
50-
Actin
C MG CQ C MG CQ C MG CQ
C
****
ns
ns
ns
ns
**
ns
NEMO remaining (NEMO/HSP90)
1.5
1.0
0.5
0.0
C MG MLN CQ C MG MLN CQ
WT
IKK DKO

Supp Figure 4

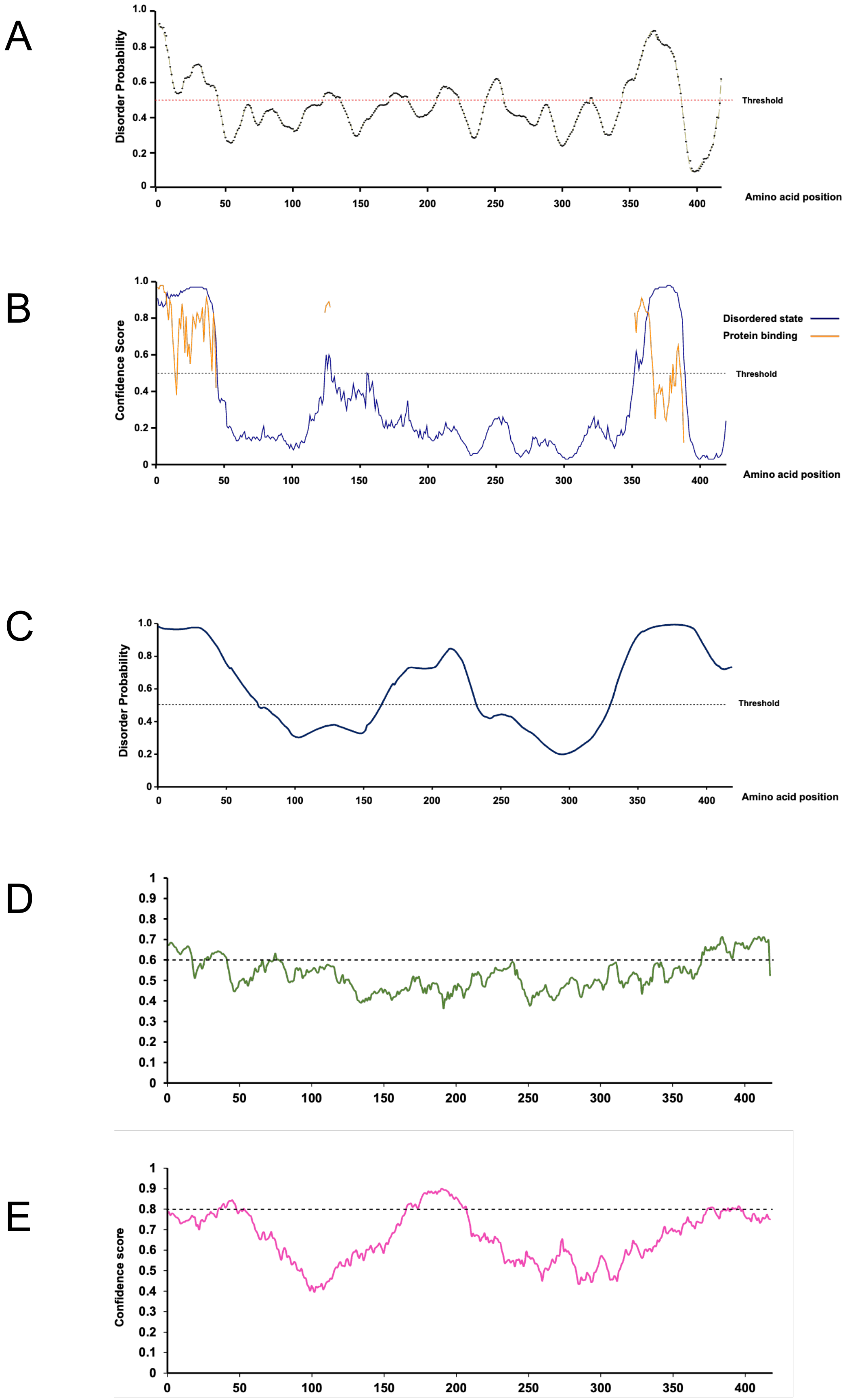

Supplementary Figure 5

A

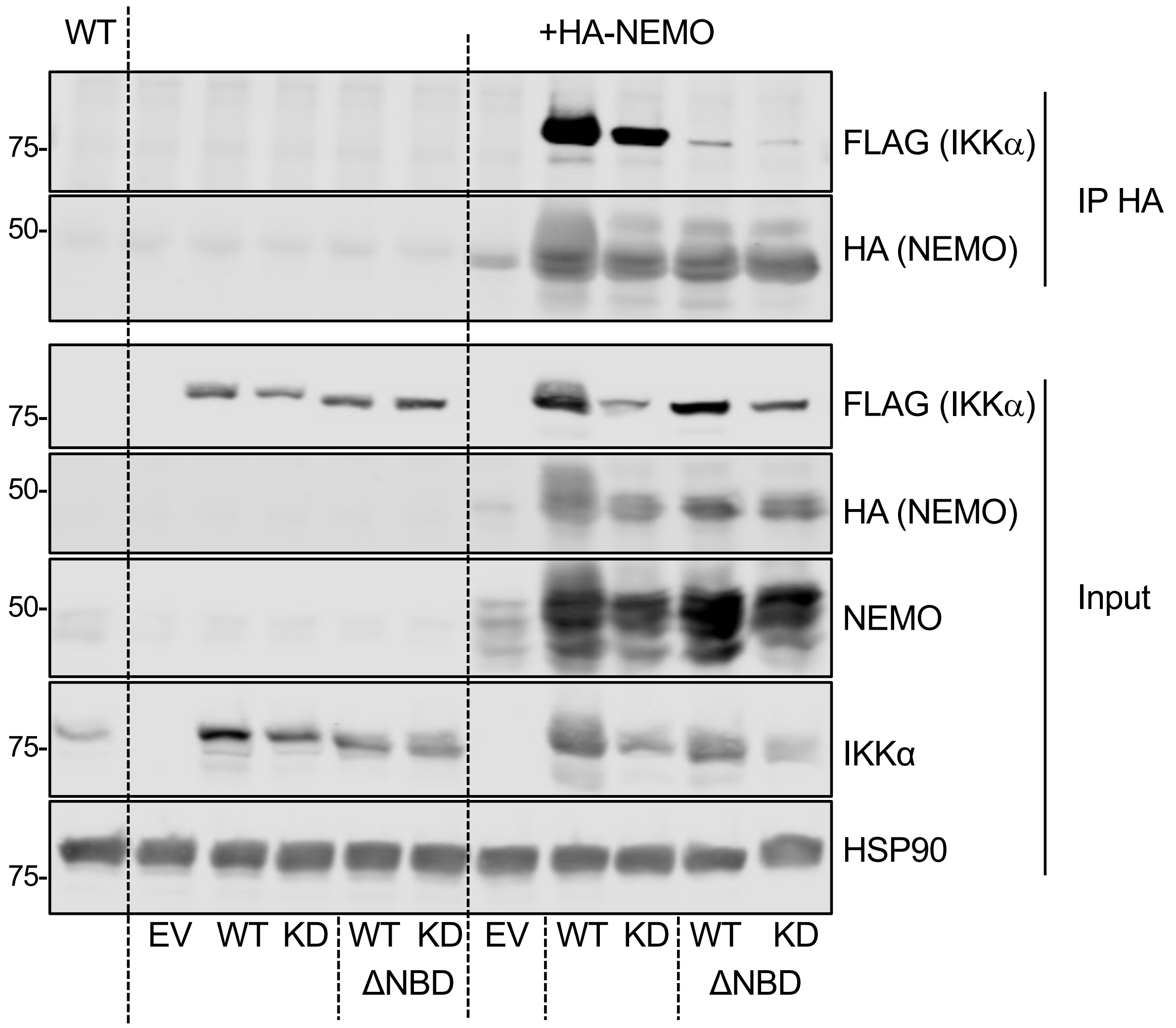


B

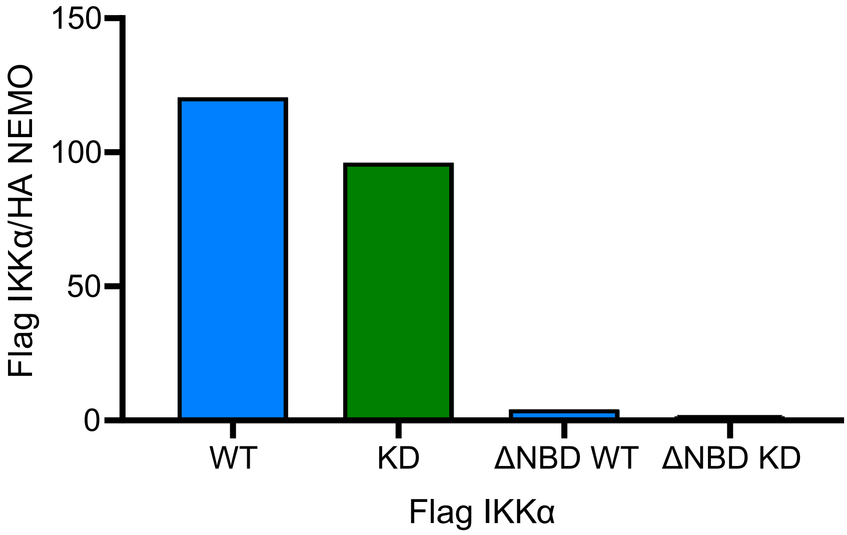
